# Tuning parameters of dimensionality reduction methods for single-cell RNA-seq analysis

**DOI:** 10.1101/2020.04.27.064816

**Authors:** Felix Raimundo, Celine Vallot, Jean Philippe Vert

**Affiliations:** Google Research, Brain team, 75009 Paris, France; CNRS UMR3244, Institut Curie, PSL Research University, 75005 Paris, France; Translational Research Department, Institut Curie, PSL Research University, 75005 Paris, France

## Abstract

**Background:** Many computational methods have been developed recently to analyze single-cell RNA-seq (scRNA-seq) data. Several benchmark studies have compared these methods on their ability for dimensionality reduction, clustering or differential analysis, often relying on default parameters. Yet given the biological diversity of scRNA-seq datasets, parameter tuning might be essential for the optimal usage of methods, and determining how to tune parameters remains an unmet need.

**Results:** Here, we propose a benchmark to assess the performance of five methods, systematically varying their tunable parameters, for dimension reduction of scRNA-seq data, a common first step to many downstream applications such as cell type identification or trajectory inference. We run a total of 1.5 million experiments to assess the influence of parameter changes on the performance of each method, and propose two strategies to automatically tune parameters for methods that need it.

**Conclusions:** We find that principal component analysis (PCA)-based methods like scran and Seurat are competitive with default parameters but do not benefit much from parameter tuning, while more complex models like ZinbWave, DCA and scVI can reach better performance but after parameter tuning.

## Introduction

Single-cell RNA sequencing (scRNA-seq) is a powerful technology to characterize the transcriptomic profile of individual cells within a population [1]. By allowing researchers to identify cell types based on their transcriptomic signatures instead of pre-defined markers, it is rapidly establishing itself as a standard tool to answer a variety of biological questions, ranging from characterizing the heterogeneity of complex tissues [2, 3] to discovering new cell types [4] or elucidating cell differentiation processes [5].

The analysis of scRNA-seq data raises, however, a number of challenges. Due to the small amount of RNA available in each individual cell, and to the technical difficulty to analyze thousands (or millions) of cells in parallel, raw scRNA-seq data have been found to be subject to a number of biases including low sequencing depth, over-dispersion and zero inflation of read counts, or sensitivity to batch effects [6, 7]. Many computational methods have therefore been developed in recent years to take into account the specificities of scRNA-seq data and address the issues of data normalization, cell type identification, differential gene expression analysis, cell hierarchy reconstruction, or gene regulatory network inference (see [8, 9] for recent reviews). In order to help practitioners choose an analysis pipeline among the many available, several studies have benchmarked algorithms and softwares for applications such as dimensionality reduction [10], clustering [11, 12], differential expression [13], or trajectory inference [14].

One shared caveat by these benchmarking efforts, however, is that the methods tested are run with their default parameters. This may not reflect what an educated user would do in practice, and does not address the practical questions of (i) whether parameter tuning is relevant at all for a given method, and (ii) how to tune parameters if needed. Recently, [15] highlighted the relevance of these questions, showing that variation autoencoders (VAE) algorithms for dimension reduction (DR) of scRNA-seq work very well once properly tuned, but are extremely sensitive to changes in parameters and can dramatically fail if not properly tuned.

Here, we propose to challenge this issue of parameter tuning focusing on methods for DR of scRNA-seq data, not only because they can be directly useful for visualization purpose, but also because DR is a common first step for most downstream applications such as cell type identification or trajectory inference [8, 9]. We propose a new benchmark protocol for DR methods, composed of ten scRNA-seq datasets of various complexity mixing experimentally characterized populations of cells, where we measure the quality of a DR method by its ability to map cells of a given cell type near each other in the representation space. Using this protocol, we benchmark five popular and representative DR methods, combining both PCA-based methods, a matrix factorization method, and VAE methods, systematically varying their tunable parameters. The resulting ∼1.5 million experiments reveal not only the performance of DR methods using their default parameters, but also their performance if parameters are properly tuned. We find in particular that principal component analysis (PCA)-based methods like scran [16] and Seurat [17] are competitive with default parameters but do not benefit much from parameter tuning, while more complex models like ZinbWave [18], DCA [19], and scVI [20] can reach better performance but after parameter tuning. We propose and evaluate two strategies to tune parameters automatically, either by changing the default parameters or by optimizing a heuristic on each new dataset. In spite of promising results for some of the methods like ZinbWave, both strategies sometimes identify very suboptimal parameters, suggesting that parameter tuning for complex DR models on dataset without ground truth annotation remains an important but largely open problem.

## Results

### A benchmark of DR methods for scRNA-seq data

A DR method takes a scRNA-seq dataset as input and maps each individual cell to a point in *d*-dimensional *representation space*, where downstream applications such as cell type prediction or lineage reconstruction are performed. In order to empirically assess the quality of DR methods and the influence of parameter tuning, we propose a benchmark protocol, summarized in Figure 1, where we collected ten diverse scRNA-seq datasets with experimentally validated cell types, and evaluate five representative DR methods, tested on a large parameter sweep, according to their ability to map cells of a given origin near to other cells of the same cell type.

**Figure 1:**
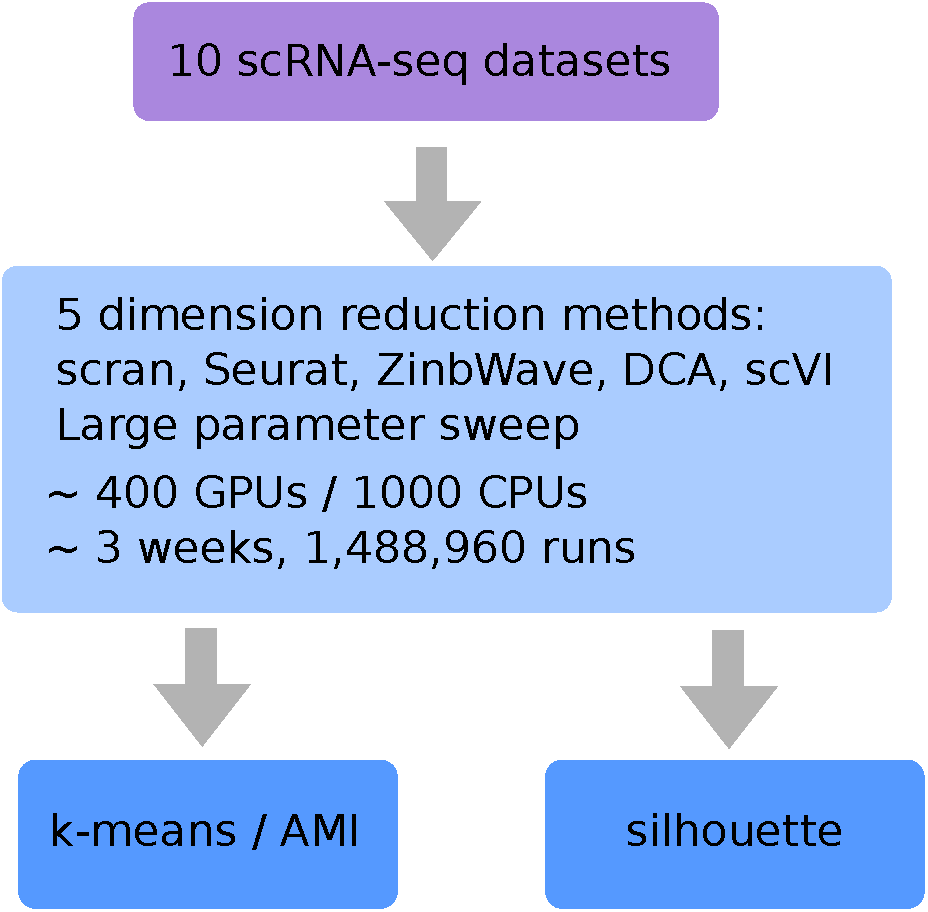
Overview of the benchmark protocol. We ran five representative DR methods, systematically varying their parameters on a large grid of values, on ten scRNA-seq datasets with known cell identity. We evaluate their ability to map cells of a given identity near other cells of the same identity, as measured by the silhouette score and the AMI after *k*-means clustering in the representation space.

Table 1 and Figure 2 summarize the main features of the ten datasets. Each dataset contains hundreds to thousands of experimentally characterized cell types, either derived from known cell lines or purified by FACS. The ten datasets vary in the technology used (10x, CEL-Seq2 or Smart-Seq2), the organism of origin (human or murine), and the overall biological complexity of the mixture. More precisely, the first five datasets (Zhengmix4eq to Zhengmixun8eq) are *in silico* mixtures of FACS purified human immune cell populations from [21] produced with 10x, comprising either equal mixes of four, five and eight cell populations, or unequal mixes of four and eight cell populations. The datasets with five and eight cell populations are particularly challenging, since they both contain five closely related T-cell populations. The next four datasets (sc_10x to sc_celseq2_5cl) are *in vivo* mixtures of three or five human cell lines from [12], sequenced by CEL-Seq2 or 10x. Finally, the last dataset is an *in silico* mixture of four FACS purified mouse tissues from [22] produced with Smart-Seq2, where we selected tissues with no overlap in cell types.

**Table 1:** Benchmark datasets. The first five datasets are derived from [21] and [11], and contain CD19+ B cells (B), CD14+ monocytes (Monocytes), CD4+ helper T cells (hT), CD56+ natural killer cells (NK), CD4+/CD45RO+ Memory T Cells (mT), CD8+/CD45RA+ Naive Cytotoxic T Cells (cytoT), CD4+/CD45RA+/CD25-Naive T cells (nT), and CD4+/CD25+ Regulatory T Cells (rT). The four next are from [12] and contain the five following cell lines: A549, H1975, H2228, H838, and HCC827. The last one is from [22].

| Dataset | Technology | Organism | N cells | Cell types | N types | Ref. |
| --- | --- | --- | --- | --- | --- | --- |
| Zhengmix4eq | 10x | Human | 3,909 | B (24.5%), Monocytes (24.5%), cytoT (25.5 %), rT (25.5 %) | 4 | [21] [11] |
| Zhengmix4uneq | 10x | Human | 6,345 | B (15%), Monocytes (30%), cytoT (8 %), rT (46 %) | 4 | [21] [11] |
| Zhengmix5eq | 10x | Human | 4,876 | hT (20%), mT (20%), cytoT (20%), nT (20%), rT (20%) | 5 | [21] |
| Zhengmix8eq | 10x | Human | 3,908 | B (12.5 %), Monocytes (14.5 %), hT (10 %), NK (15 %), mT (12.5 %), cytoT (10 %), nT (13 %), rT (12.5 %) | 8 | [21] [11] |
| Zhengmix8uneq | 10x | Human | 6,350 | B (7.5%), Monocytes (15%), hT (8%), NK (4%), mT (15%), cytoT (4%), nT (23%), rT (4%) | 8 | [21] |
| sc_10x | 10x | Human | 902 | H1975 (34.5 %), H2228 (35 %), HCC827 (30.5 %) | 3 | [12] |
| sc_10x_5cl | 10x | Human | 3,918 | A549 (32 %), H1975 (11 %), H2228 (19.5 %), H838 (22.5 %), HCC827 (15 %) | 5 | [12] |
| sc_celseq2 | CEL-Seq2 | Human | 274 | H1975 (41 %), H2228 (29.5 %), HCC827 (29.5 %) | 3 | [12] |
| sc_celseq2_5cl | CEL-Seq2 | Human | 895 | A549 (36 %), H1975 (14.5 %), H2228 (14 %), H838 (22 %), HCC827 (13.5 %) | 5 | [12] |
| TabulaMuris | Smart-Seq2 | Murine | 12,081 | Brain (36.5 %), Intestine (31.5 %), Skin (19 %), Spleen (13 %) | 4 | [22] |

**Figure 2:**
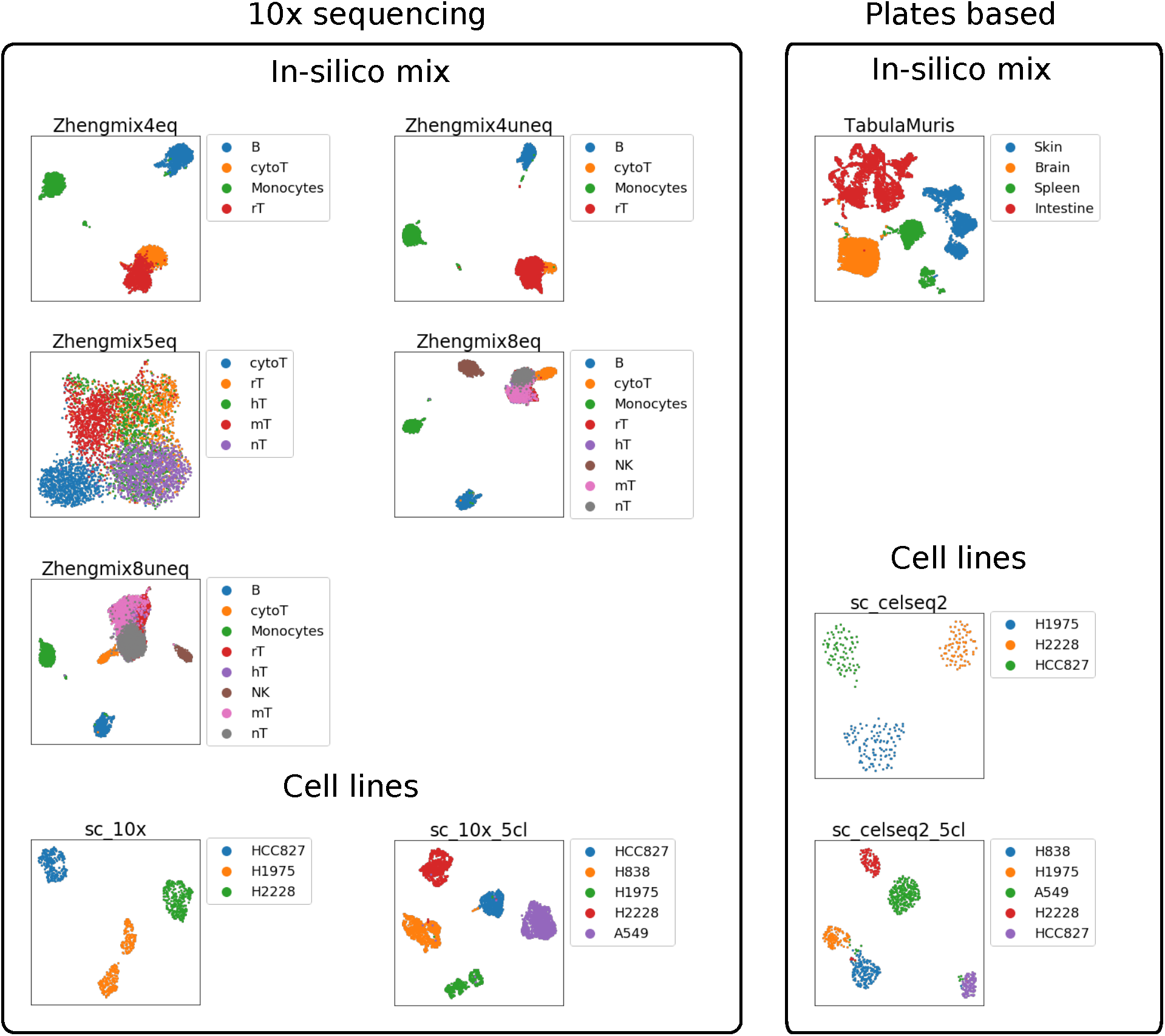
UMAP representation of the ten scRNA-seq datasets, run after processing of the count matrices with Seurat with default parameters.

We use this benchmark to evaluate the performance of five popular computational pipelines for DR of scRNA-seq data: scran [16], Seurat [17], ZinbWave [18], DCA [19], and scVI [20]. These pipelines are all publicly available as R or Python packages, and can process datasets containing thousands of cells in a reasonable amount of time (less than 12 hours on a GPU/CPU with 10 cores). They all implement processing steps speficic to scRNA-seq data together with representative DR methods including principal component analysis (PCA) for scran and Seurat, matrix factorization for ZinbWave, and (variational) autoencoders for DCA and scVI. While these pipelines also implement various downstream tasks such as cell clustering or differential expression analysis, we restrict our analysis to the DR step.

To quantify the ability of a DR method to map biologically similar cells to similar locations in the representation space, we use two complementary measures: the silhouette, on the one hand, and the adjusted mutual information (AMI) when the cells are clustered with the *k*-means algorithm, on the other hand (see details in Material and Methods). Both measures vary between 0 for a random embedding to 1 for an embedding that perfectly preserves the cell type information. Silhouette is a measure agnostic to any particular clustering algorithm, and measures how close a cell is to other cells of the same type compared to cells of different types; for the silhouette to be large, cell types must not only be separated, but also form compact clusters far from each other. AMI, on the other hand, directly measures how well a particular clustering algorithm recovers known cell types, and is therefore a good proxy for the performance of cell type identification as a downstream task of DR. Both measures are frequently used to assess the performance of cell embedding techniques [18, 11, 20, 23, 10]. Another standard measure to assess the ability of a clustering algorithm to recover known classes is the adjusted rand index (ARI) [11, 20, 23, 10]; however, we found that AMI and ARI are extremely correlated (Additional file 1: Fig. S1) and lead to similar conclusions, so we just report results based on AMI below.

### Performance of five popular DR methods with default parameters

We first assess the performance of each method with its default parameters, except for the dimension of the representation space which we arbitrarily set to 10 for all methods. Indeed, the performance scores (AMI and silhouette) strongly vary with the dimension (Additional file 1: Fig. S2 and S3), so fixing the dimension allows to compare more fairly the different DR methods. Figure 3.A (with “default” legend) shows the performance reached by each method on each dataset, in terms of AMI (left) and silhouette (right), and Table 2 summarizes the mean performance reached by each method over the datasets.

**Table 2:** Mean performance on the ten datasets of each method in terms of AMI and silhouette. The “Default” columns correspond to the performance of each method using its default parameters, with a dimension of 10. The “Best” column corresponds to the best performance reached after varying the parameters. The “ANOVA AMI heuristic” column corresponds to the performances of the new default parameters described in section. The “silhouette heuristic” column corresponds to the performance of the heuristic described in section

| Method | Mean AMI |  |  |  | Mean silhouette |  |  |  |
| --- | --- | --- | --- | --- | --- | --- | --- | --- |
|  | Default | Best | ANOVA AMI heuristic | silhouette heuristic | Default | Best | ANOVA AMI heuristic | silhouette heuristic |
| scran | **0.840** | 0.868 | 0.841 | 0.741 | 0.362 | 0.547 | 0.396 | 0.494 |
| Seurat | 0.788 | 0.860 | 0.814 | 0.683 | 0.369 | 0.543 | 0.490 | 0.373 |
| ZinbWave | 0.780 | **0.896** | **0.851** | **0.825** | 0.249 | 0.609 | **0.562** | **0.591** |
| DCA | 0.758 | 0.885 | 0.837 | 0.583 | **0.396** | **0.639** | 0.403 | 0.381 |
| scVI | 0.560 | 0.872 | 0.709 | 0.510 | 0.384 | 0.621 | 0.482 | 0.318 |

**Figure 3:**
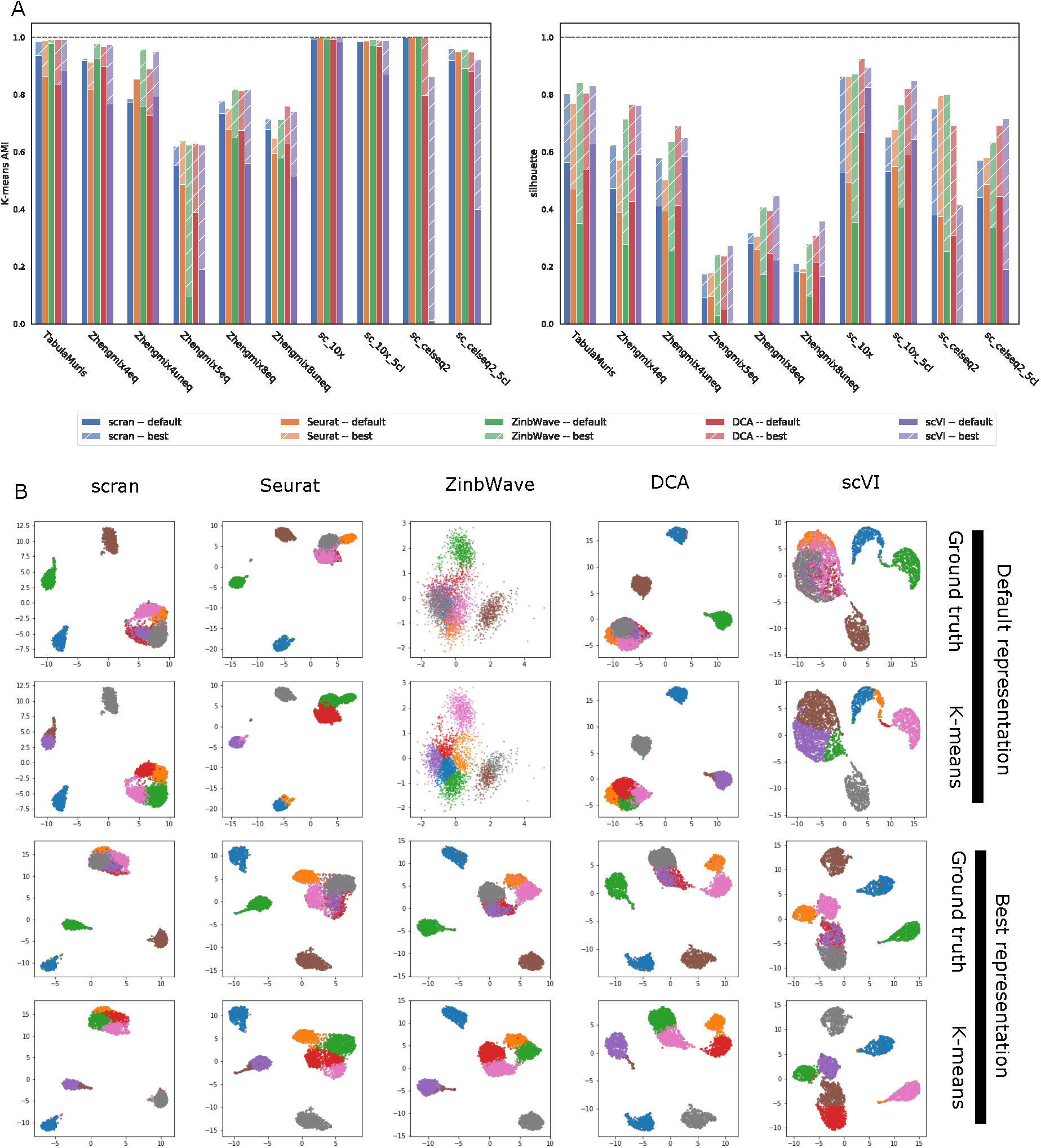
Performance of five DR pipelines (scran, Seurat, ZinbWave, DCA and scVI) with default parameters and a dimension of 10 (legend “default”) or after parameter optimization (legend “best”) on our benchmark of ten datasets. **A**. AMI (left) and silhouette (right) reached by each method on each dataset. **B**. UMAP representation of Zhengmix8eq after DR by each method (in column) using default parameters (top two rows) of after parameter optimization (bottom two rows). In each row, cells are colored either based on their true cell type (rows 1 and 3) or based on a *k*-means clustering.

As expected, Figure 3.A clearly shows that for all methods, the performance varies across datasets, in a rather consistent manner. In terms of AMI, the four cell lines datasets tend to be the easiest (AMI>0.9 for most methods), followed by TabulaMuris and the two Zhengmix mixtures of four cell lines (AMI in the range 0.7 ∼ 0.9 for most methods), followed by the three Zhengmix mixtures containing the five closely related T-cell populations (AMI in the range 0.1 ∼ 0.7 for most methods). The silhouette scores overall follow the same trend, although the difference between the first two groups of datasets is less pronounced. Interestingly, we see that in cases where several DR methods allow to almost perfectly cluster the cell types, such as sc_10x or sc_10x_5cl, the silhouette is usually far from 1 and varies across methods, illustrating the complementarity of both measures.

Besides variations across datasets, we also observe variations across DR methods. As shown in Table 2, scran has the best AMI on average (mean AMI=0.84), followed by Seurat, ZinbWave and DCA (mean AMI=0.75 ∼0.79), but this ranking is not statistically significant (p-value > 0.05 for Wilcoxon one-way test), while scVI is clearly behind (mean AMI=0.56) (p-value < 0.05 for all methods). As suggested in Figure 3.B on Zhengmix8eq, for example, scVI with default parameters does not manage to clearly isolate the three non-T cell clusters, resulting in errors in *k*-means clustering. In terms of silhouette, all methods are very similar (mean silhouette=0.36 ∼ 0.39), except for ZinbWave which is clearly behind (p-value < 0.05 for all methods). The reason why ZinbWave tends to have a correct AMI but a poor silhouette is suggested by Figure 3.B, where we see on Zhengmix8eq that ZinbWave (with default parameters) produces a representation good enough for *k*-means to correctly recover most of the cell types, but where the different clusters look much less compact and separated from each other than with other DR methods. The p-values for the comparisons can be found in Additional file 1: Fig. S16-S17.

While these relative performances hold for a representation in dimension 10, the performance of some methods fluctuates with the dimension of the embedding space (Additional file 1: Fig. S2 and S3). While scran and Seurat are rather insensitive to increase in dimension after 10, DCA’s mean AMI tends to increase in higher dimensions, while ZinbWave’s and scVI’s mean AMI decrease with the dimension, suggesting that different methods need more or less dimension to capture the same biology.

This average performance of DR methods hides important variations across datasets, as visualized in Figure 3.A. For example, we see that scVI has specifically poor performance compared to other methods on the two CEL-Seq2 datasets, which may be due to a particularity of this technology or to the fact that both datasets (sc_celseq2 and sc_celseq2_5cl) have a relatively small number of cells. The difference in AMI across methods, and the good behavior of the linear models underlying scran, Seurat and ZinbWave, is most visible on the “difficult” Zhengmix mixtures containing the five closely related T-cell populations, while the difference in silhouette, and the good behaviour of the nonlinear models underlying DCA and scVI, is more visible in the “easy” datasets where all methods have an AMI above 0.7.

### Performance reachable across a parameter sweep

While using a computational pipeline with default parameters is often the method of choice for practitioners, there is little guarantee that default parameters are adapted to all situations. In particular, the performance of deep learning-based methods for scRNA-seq analysis was shown to be highly sensitive to choices of parameters [15]. In order to assess the performance of each DR method if all parameters were properly tuned in a dataset-specific way, we now run each method by sweeping all tunable parameters across a large grid of values, as summarized in Table 3, and compute the performance reached by each method after cherry picking *a posteriori* the best parameters. Note that the resulting performance is therefore an upper bound on the performance that each method can reached if parameters are tuned without knowing the ground truth.

**Table 3:**
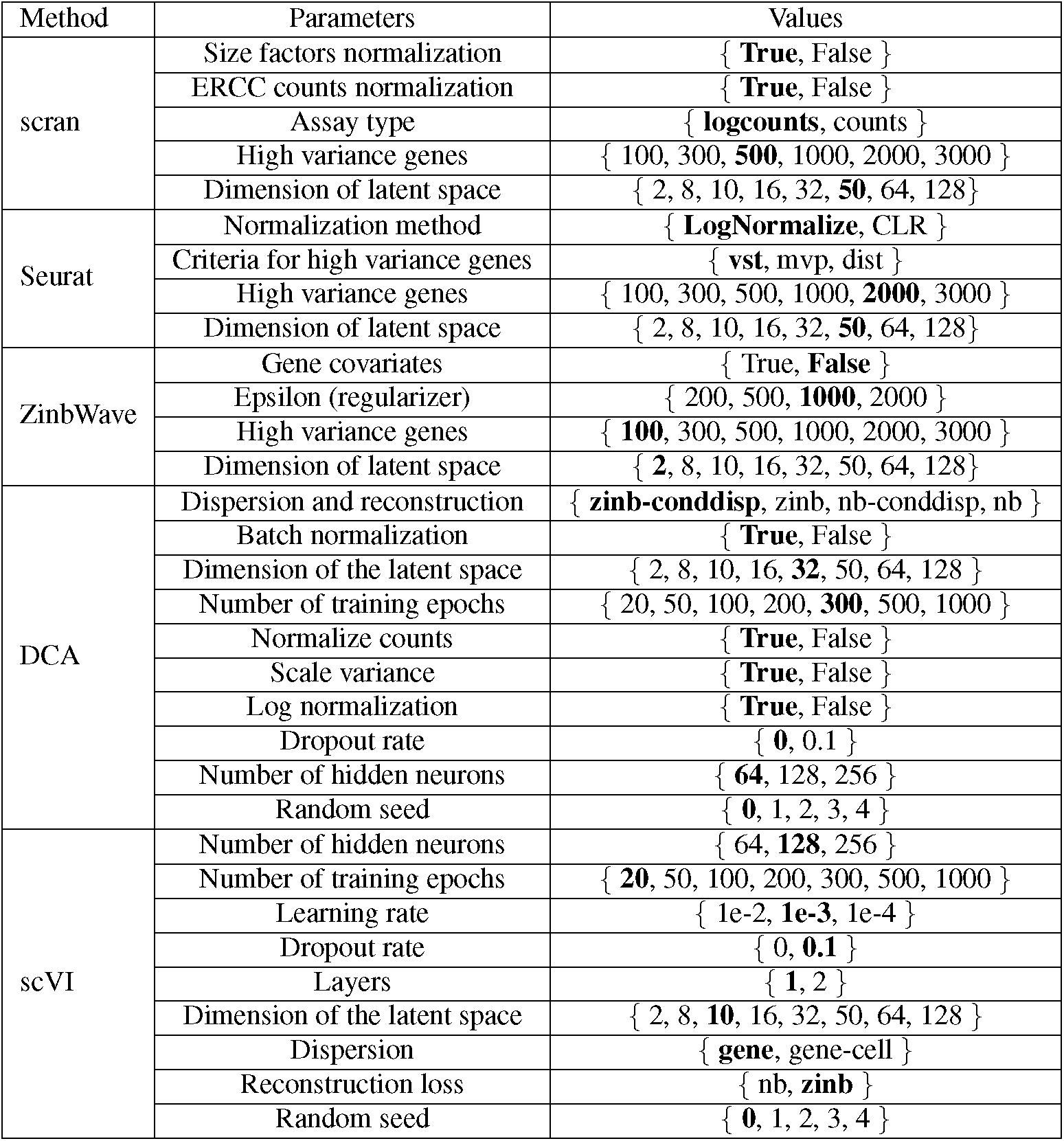
Description of parameter sweep. For each method (first column), we vary a number of tuneable parameters (second column) systematically over a grid of values (third column). The bold value in the third column is the default value.

| Method | Parameters | Values |
| --- | --- | --- |
| scran | Size factors normalization | { **True**, False } |
|  | ERCC counts normalization | { **True**, False } |
|  | Assay type | { **logcounts**, counts } |
|  | High variance genes | { 100, 300, **500**, 1000, 2000, 3000 } |
|  | Dimension of latent space | { 2, 8, 10, 16, 32, **50**, 64, 128 } |
| Seurat | Normalization method | { **LogNormalize**, CLR } |
|  | Criteria for high variance genes | { **vst**, mvp, dist } |
|  | High variance genes | { 100, 300, 500, 1000, **2000**, 3000 } |
|  | Dimension of latent space | { 2, 8, 10, 16, 32, **50**, 64, 128 } |
| ZinbWave | Gene covariates | { True, **False** } |
|  | Epsilon (regularizer) | { 200, 500, **1000**, 2000 } |
|  | High variance genes | { **100**, 300, 500, 1000, 2000, 3000 } |
|  | Dimension of latent space | { **2**, 8, 10, 16, 32, 50, 64, 128 } |
| DCA | Dispersion and reconstruction | { **zinb-conddisp**, zinb, nb-conddisp, nb } |
|  | Batch normalization | { **True**, False } |
|  | Dimension of the latent space | { 2, 8, 10, 16, **32**, 50, 64, 128 } |
|  | Number of training epochs | { 20, 50, 100, 200, **300**, 500, 1000 } |
|  | Normalize counts | { **True**, False } |
|  | Scale variance | { **True**, False } |
|  | Log normalization | { **True**, False } |
|  | Dropout rate | { **0**, 0.1 } |
|  | Number of hidden neurons | { **64**, 128, 256 } |
|  | Random seed | { **0**, 1, 2, 3, 4 } |
| scVI | Number of hidden neurons | { 64, **128**, 256 } |
|  | Number of training epochs | { **20**, 50, 100, 200, 300, 500, 1000 } |
|  | Learning rate | { 1e-2, **1e-3**, 1e-4 } |
|  | Dropout rate | { 0, **0.1** } |
|  | Layers | { **1**, 2 } |
|  | Dimension of the latent space | { 2, 8, **10**, 16, 32, 50, 64, 128 } |
|  | Dispersion | { **gene**, gene-cell } |
|  | Reconstruction loss | { nb, **zinb** } |
|  | Random seed | { **0**, 1, 2, 3, 4 } |

In total, sweeping across the grid of parameters results in 384 different runs per dataset for scran and ZinbWave, 288 for Seurat, 40,320 for scVI, and 107,520 for DCA, hence a total of 1,488,960 DR experiments. Running all experiments took several weeks on a dedicated cluster of 1,000 CPUs and 400 GPUs. Out of these runs 60% ran correctly for scran, 99.97% for Seurat, 96.51% for ZinbWave, 97.96% for DCA, and 100% for scVI. The low number of runs for scran is mostly due to the absence of ERCC in all the 10X datasets, and because the size factor computation on TabulaMuris failed. The failures for ZinbWave and DCA were due to memory issues (either of the GPU or CPU). There was a single failure for Seurat whose cause has not been identified.

We report in Figure 3.A and Table 2 (with “Best” legend) the best value reached across the parameter sweep on each dataset, in addition to the performance reached with the default parameters. Overall, we see that for all methods, a gain can result from parameter tuning compared to using default parameters. For AMI, the mean gain across datasets ranges from 0.311 for scVI to 0.028 for scran, while for for silhouette, it ranges from 0.185 for scran to 0.360 for ZinbWave. Seurat and scran are the methods that benefit least from parameter tuning, suggesting that default parameters are already good choices across most datasets. Autoencoder-based DCA and scVI benefit more for parameter tuning, and outperform scran and Seurat in mean AMI after parameter tuning, confirming the importance of parameter tuning for these models [15]. Since the number of parameters tested for these models is also two orders of magnitude larger than for Seurat, scran and ZinbWave, the “best” performance after cherry-picking the best parameters may be over-optimistic for DCA and scVI. As for ZinbWave, a good choice of parameters leads to the best mean AMI across methods (0.896), and the largest improvement in silhouette compared to default parameters.

More precisely, for all cell line datasets and for TabulaMuris, parameter tuning allows all method to reach an almost perfect AMI, including methods like scVI that have a very poor performance with default parameters on sc_celseq2 and sc_celseq2_5cl. On the same datasets, parameter tuning brings an important improvement to the silhouette score of 0.2 to 0.6 to all methods. After parameter tuning, both encoder-based methods (DCA and scVI) tend to outperform ZinbWave, which tends to outperform both PCA-based methods scran and Seurat in terms of silhouette. This highlights the possibilities of nonlinear DR methods to perform DR even on simple datasets, but the need to correctly tune parameters in order to reveal their full potential.

On the immune cell datasets, we see again that tuning parameters allows to boost performance and bridge important gaps between methods in terms of silhouette, and that after parameter optimization both autoencoder-based methods slightly outperform ZinbWave, which slightly outperforms both PCA-based methods. The AMI performance of all methods is also improved by parameter optimization for all methods but scran, and we see no clear and consistent winner after parameter optimization. This suggests that simple PCA-based methods like scran, even with default parameters, are good enough to match the performance of more complex models after parameter tuning in terms of AMI; however the better silhouette of more complex models once tuned may be an advantage for other downstream applications beyond clustering by *k*-means. The benefits of parameter tuning is further illustrated in Figure 3.B, which shows a UMAP visualization of the representation space learned by the different methods with the default parameters (bottom two rows) or after parameter tuning for AMI (top two rows), for the Zhengmix8eq dataset. For ZinbWave and scVI, which strongly benefit from parameter tuning in this case, we see that the dataset looks very different with the default parameters or after parameter optimization, the different cell types being but better separated in the later case.

Regarding the absolute performance reached across the datasets, it is interesting to note that the best AMI scores for Zhengmix5eq and Zhengmix8uneq (the two hardest datasets) are only between 0.6 and 0.7, when they are above 0.9 for Zhengmix4eq (the easiest one). This suggests that if current DR methods are good at identifying sufficiently different cell types, they have difficulties to differentiate very similar cell types, a variety of T cells in our case, even after parameter optimization.

To check how the AMI results are influenced by the choice of the clustering algorithm (here *k*-means), we also computed the AMI performance when cells are clustered by Ward’s hierarchical clustering (HC) or the Louvain algorithm [24]. As shown on Additional file 1: Fig. S4, the performance barely changes when *k*-means is replaced by HC. When Louvain clustering is used, the overall performance decreases on most datasets, possibly because we used default parameters for this algorithm, while *k*-means and HC are run knowing in advance the correct number of clusters. The relative ordering of the different method is overall conserved, suggesting our conclusions based on AMI are robust to the clustering method.

### Influence of parameters on performance

Having shown that parameter tuning has the potential to boost the performance of all methods for all datasets compared to using default parameters, we now investigate in more details the influence of each parameter on the performance. For that purpose, we estimate the mean contribution of each parameter value on the performance (AMI or silhouette) with a factorial analysis of variance (ANOVA, see Methods) procedure. We find that all parameters of all methods, except for “gene covariate” for ZinbWave on the silhouette, have a significant influence on both AMI and silhouette (ANOVA p-value < 0.05), and summarize in Additional file 1: Table S1 and S2 the potential effect of tuning each parameter by comparing the best and worse contributions to performance among the values it can take. We see that some parameters can have a very important effect, such as proper log normalization in scran which on average can boost the AMI by 0.50, or the choice of dimension in the latent space for ZinbWave that can boost the silhouette by 0.56 on average. If we arbitrarily define a parameter as “influential” if its potential effect is more than 0.05 on AMI or silhouette, we see that in addition to the dimension of the representation space which is influential for all methods, scran, Seurat and ZinbWave have one influential parameters (log normalization for scran; normalization method for Seurat; number of top genes for ZinbWave), DCA has two (batch normalization and normalize counts), and scVI has three (dispersion, number of training epochs and learning rate), suggesting that more care in parameter tuning is needed for the autoencoder-based methods than for the matrix factorization-based methods.

As shown in Table 3 and Additional file 1: Table S1, the best parameter values in terms of mean contribution to the performance are not always the default parameters of each method. This suggests that the best parameter values identified by our analysis, which we refer to below as “ANOVA AMI heuristic” when we pick the parameter values that have the largest positive influence on AMI, may be interesting to use as new default parameters for each method. Note that, by construction, the ANOVA AMI heuristic parameters are the same for all datasets. To test this hypothesis, we report the performance of each DR method using the ANOVA AMI heuristic as default parameters in Additional file 1: Fig. S5, and summarize the mean performance across datasets in Table 2.

We see that, on average, all methods benefit from the ANOVA AMI heuristic compared to the existing default parameters, particularly scVI, DCA and ZinbWave in terms of AMI, and particularly ZinbWave, Seurat and scVI in terms of silhouette. In particular, ZinbWave outperforms all other methods, both in AMI and in silhouette, with the ANOVA AMI heuristic. Interestingly, we see in Additional file 1: Fig. S5 that for all methods, the AMI increases on almost all datasets with the ANOVA AMI heuristic. Of course these promising results should be taken with care, given that we evaluate the performance of the ANOVA AMI heuristic on the datasets used to perform the ANOVA, but they suggest a systematic approach for method developers to set default parameters.

We now investigate in more details to what extent further performance gain may result from parameter tuning on each dataset separately, as opposed to setting new default parameter values common to all datasets. For that purpose we estimate again the contribution of each parameter in the final AMI and silhouette, allowing a different contribution in different datasets by adding interaction terms in the factorial ANOVA model between the dataset, on the one hand, and the tunable parameters, on the other hand (see Methods). Note that we remove from this analysis a few parameters that were not tested on all datasets: ERCC for scran on 10x datasets, and sum factor normalization for scran on TabulaMuris. Additional file 1: Fig. S6-S15 summarize the contributions of each parameter in each dataset for AMI and silhouette, as estimated from the factorial ANOVA with interactions. All interactions between dataset and parameters are significantly non zero (p-value < 0.05), except for the interaction between dataset and the “gene covariate” parameter of ZinbWave, showing that the influence of most parameters on the final performance is not the same across datasets. To assess whether dataset-specific parameter tuning is useful, we now check for each parameter whether the default value provided by the ANOVA AMI heuristic, which identifies the best values on average, is also the best or within 0.05 of the best for both AMI and silhouette on *all* datasets. Based on this criterion, we find that for the AMI scran, Seurat and ZinbWave have one parameter that benefits from dataset-dependent tuning (number of top genes for scran and Seurat; dimension of the latent space for ZinbWave), DCA has two (scale variance and dropout rate), and scVI has three (dimension of the latent space, number of hidden neurons and number of training epochs); and for the silhouette, scran, Seurat, ZinbWave and scVI have one parameter that benefits from dataset-dependent tuning (dimension of the latent space for scran and Seurat; number of top genes for ZinbWave; number of training epochs for scVI), and DCA has two (scale variance and dropout rate). Additional file 1: Table S1 and Table S2 detail the potential gain in dataset-specific tuning for each parameter of each method. This therefore confirms the potential benefit of tuning parameters on each dataset, particularly for autoencoder-based methods.

### Tuning parameters in practice

Having shown that DR methods benefit from various degrees of parameter tuning, we now discuss the question of how this can be done in practice. Indeed, our strategy so far to identify the best parameters and evaluate their influence on performance is only possible when one knows the true cell type for each cell in the population, but such an oracle is usually not available in practice. In the absence of such information, one must therefore rely on quantitative heuristics or qualitative validation by domain experts, e.g., by looking at the distribution of cells in the representation space and assessing whether it shows some promising structure such as clusters.

As a first step towards an automated way to tune parameters in a dataset-specific way, we now propose a simple quantitative and objective heuristic to tune parameters, which we call the *silhouette heuristic*, and evaluate its performance on our benchmark. The silhouette heuristic measures how well a distribution of cells in the representation space looks like a possible clustering of distinct cell types. Given a set of cells in a representation space, typically a dataset of cells processed by a DR method with some parameter values, the silhouette heuristic first runs a *k*-means clustering algorithm on the cells in the representation space, and then computes the silhouette score of the dataset with respect to the cell types assigned by the *k*-means clustering. In particular, if the *k*-means clustering identifies the true cell types, then the silhouette heuristic boils down to the silhouette score with respect to the true cell types. To tune parameters for a DR method on a dataset, we then just compute the silhouette heuristic over a grid of candidate parameter choices, and select the values that maximize it. Here we chose *k* to be the true number of cell populations, which we already know in advance. In real applications that number may not be known and has to be estimated with prior knowledge or other heuristics. Thus our heuristic here only works if the practitioner already knows the true number of populations.

Additional file 1: Fig. S5 shows the performance all methods on all datasets when parameters are tuned by maximizing the silhouette heuristic, and Table 2 summarizes the mean performance across datasets. The UMAP projection of the resulting DRs can be seen in Figure 4.

**Figure 4:**
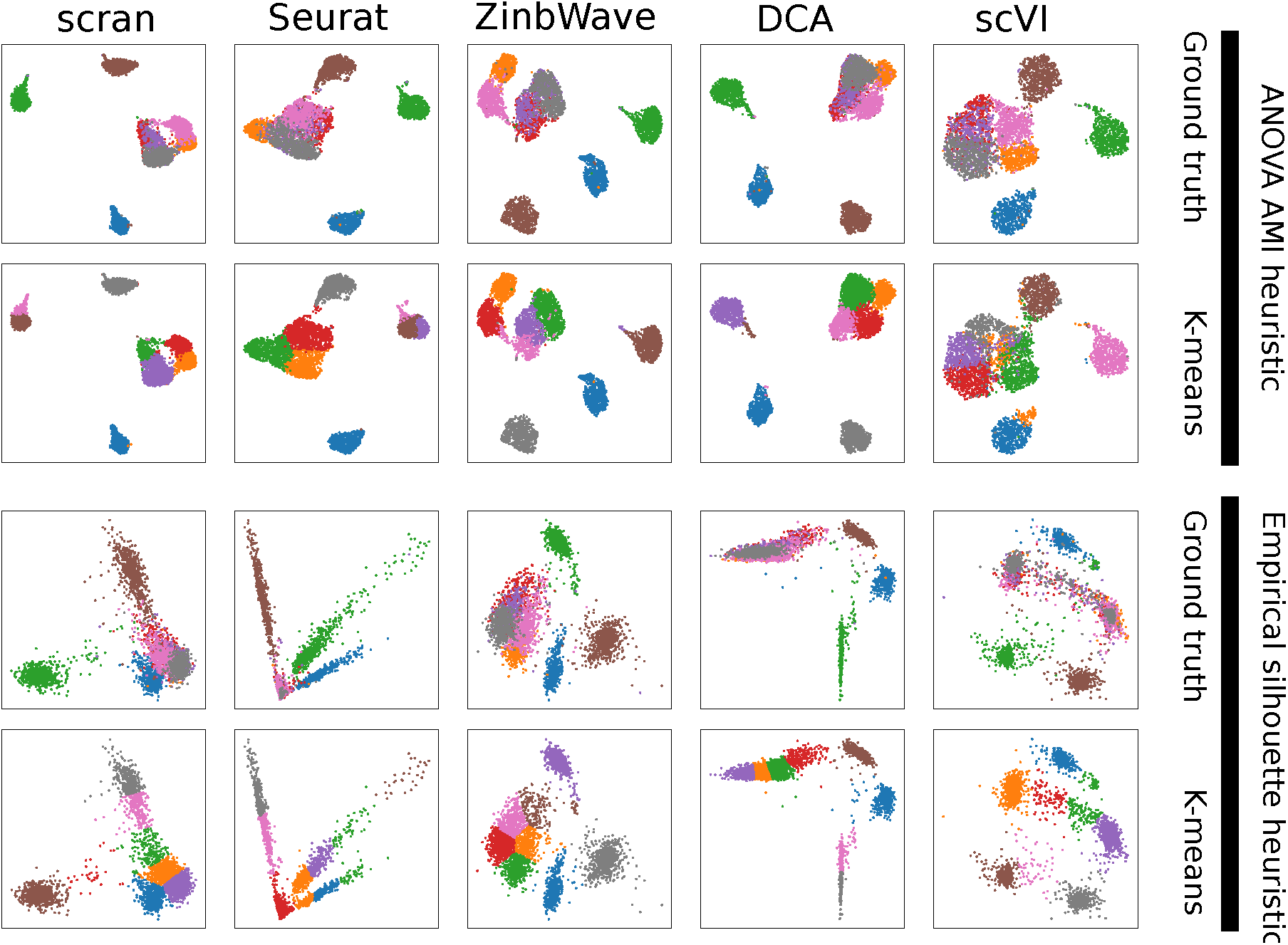
UMAP representation of Zhengmix8eq after DR by each method (in column) using the ANOVA AMI (top two rows) or empirical silhouette (bottom two rows) heuristic to tune parameters. In each row, cells are colored either based on their true cell type (rows 1 and 3) or based on a *k*-means clustering.

We can see that the silhouette heuristic works very well for ZinbWave, where it always identifies parameters equal or very close to the best ones in terms of silhouette, and is comparable to the default parameters in terms of AMI. It also works well for all methods for simple datasets like sc_10x, sc_10x_5cl and sc_celseq2_5cl, where it also identifies parameters equal or close to the best ones for the silhouette score. As shown by the good AMI performance, these are cases where the initial *k*-means clustering recovers the correct clustering with good accuracy. However, there are also cases of surprising failures on easy datasets, for example for DCA on TabulaMuris, where the parameter set selected by the silhouette heuristic has a very bad AMI and silhouette with respect to the true labels, probably because the initial clustering selected by the silhouette heuristic completely fails to identify the cell types but nevertheless leads to a good empirical silhouette, while simply using the default parameters gives an almost perfect clustering in terms of AMI and a decent silhouette. On the more challenging immune cell datasets, on the other hand, the silhouette heuristic does not seem to be useful (except for ZinbWave). It leads to worse parameters than the default ones for all methods but ZinbWave on the difficult Zhengmix8uneq and Zhengmix5eq datasets, except for scran on the later one. For the easier Zhengmix4eq and Zhengmix4uneq, it leads to better parameters for all methods but scVI. In summary, we see that automatically tuning parameters to try to increase the silhouette using the silhouette heuristic only works well on relatively simple cases, up to possible dramatic errors, but on more challenging situations where there is no clear separation between cell types then it can lead to disastrous choices by overfitting a bad initial clustering. ZinbWave is an exception where, in our benchmark, the silhouette heuristic gives consistently good results. Proposing other heuristics that really help tune parameters is an important open challenge. In order to stimulate further research in that direction, we provide as supplementary data an original dataset consisting of 5,000 embeddings (100 for each single cell dataset and for each method, corresponding to 100 random choices of different parameters, including the default ones and the ones achieving the best AMI and silhouette).

## Discussion

In this study we have systematically compared the performances of five representative and popular DR methods over ten datasets with known experimental ground truth, representing various levels of biological complexity. Importantly, we have extensively investigated how the choice of parameters for these methods influence their performances, and discussed various ways to properly tune parameters. This can inform practitioners about both the capacity of these methods, as well as on the amount of work required to properly tune them.

When properly tuned, we did not observe huge differences in performance between the methods, particularly in terms of AMI. Both PCA-based methods (scran and Seurat) are nevertheless outperformed by ZinbWave and both autoencoder-based methods (DCA and scVI), particularly in terms of silhouette. On the other hand, we also found that autoencoder-based methods have more parameters that require careful tuning than ZinbWave and PCA-based methods. We illustrated with the silhouette heuristic that automatic parameter tuning is not always easy when the true cell types are not known, and can lead to disastrous results. Interestingly, a similar conclusion was reached by [15], who highlighted the impact of parameters choices in a variational autoencoder-based model, and the need to tune them. We benchmarked DR methods using downstream analysis-agnostic metrics, mostly for simplicity and because of a lack of ground truth, other than simulations, for tasks such as trajectory inference. It would be interesting to investigate in further studies how well our metrics translate to downstream applications: in particular which metric out of the AMI and silhouette correlates best with downstream performances.

An interesting result of our study is the large drop in performance, for all methods, on the immune cell mix data compared to the cell lines. This shows that good performances in the later does not necessarily translate to good performances in the former. In particular the performances on Zhengmix8eq and Zhengmix8uneq showed that the methods failed to properly separate the various T-cells populations, probably due to their relative similarity compared to the other cell populations present in the datasets. Being able to separate similar populations is of utmost relevance when investigating, for example, early tumor development, where the tumor cell population still displays a transcriptome very similar to that of cells of the organ of origin. For example in the case of triple negative breast cancer, in which tumor cells originate from normal luminal cell populations, it is crucial to be able to distinguish the various states the tumor cells undergoes towards full transformation, in order to properly target these cells at an early stage of the disease. Our study shows that efforts are still needed to develop methods able to robustly discover cell populations in complex mixtures.

## Material and methods

### Clustering algorithms

To perform the clustering we used 3 different algorithms: *k*-means, hierarchical clustering with Ward’s linkage, and Louvain. Both *k*-means and Ward were run with correct number of clusters, and default parameters using scikit-learn’s implementation [25], version 0.21.3. For Louvain, the nearest neighbors graph construction was done with scanpy (version 1.3.8) using default parameters, and the clustering was also run with default parameters using the ‘taynaud’ flavor. Note that since we had to automatically run Louvain on all embeddings, as done in [23], we could not properly tune the resolution nor the size of the neighborhood and thus Louvain could either overcluster or undercluster.

### Performance metrics

Given a set of cells with given ground truth labels, we consider two metrics to measure how well a mapping of those cells in a representation space fits the ground truth labels.

The first metric is the mean *silhouette*, defined as the average over all cells of each cell’s silhouette. The silhouette of a given cell *x* is defined as

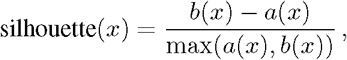

where *a*(*x*) is the average Euclidean distance between *x* and the other cells of the same class, and *b*(*x*) is the average Euclidean distance between *x* and cells in the closest different class. We used the implementation of scikit-learn.

The second metric is the AMI [26], which measures how well the ground truth clustering *U* matches the clustering *V* found by a clustering algorithm (such as *k*-means) in the representation space. The AMI is formally defined as:

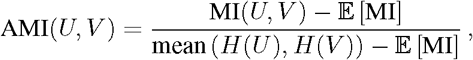

where MI(*U, V*) is the mutual information between both clustering *U* and *V, H* is the entropy function, and 𝔼 [MI] is the expected mutual information between *U* and a random clustering *V*. We used scikit-learn’s implementation with default parameters for both the *k*-means algorithm (setting *k* equal to the ground truth number of classes), and to compute the AMI.

The ARI [27] is another metric that can be used to compare how two clusterings match each other. It is formally defined as:

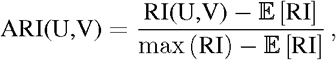

where RI(*U, V*) is the Rand index, i.e., the fraction of pairs of samples which are either in the same group or in different groups in both *U* and *V*, 𝔼 [RI] is the expected Rand index between *U* and a random clustering *V*, and max (RI) is the largest possible Rand index between *U* and any *V*. We also used scikit-learn’s implementation of the ARI, and compared the clustering obtained with *k*-means with the ground truth labels as we did for the AMI.

### Statistical analysis

To analyze results we used R (version 3.6) to perform T-tests and run ANOVA analysis with the AovSum function from the FactoMineR package (version 2.3) [28]. All parameter values were turned into factors for the ANOVA analysis. To compare the methods presented in Table 2, as shown in Additional file 1:Fig. S16-S17, we used the wilcoxon function from the scipy package [29] (version 1.14), with default parameters, except for alternative which was set to “greater” in order to have a one-way test. Note that we only had 10 samples, which is small for that test.

### Computational methods

The five DR methods were downloaded from their canonical package. We used scran version 1.14.6, Seurat version 3.1.5, ZinbWave version 1.8.0, DCA version 0.2.3, and scVI version 0.4.0. We followed either the tutorials or vignettes available at the time of download for each methods to use them. The selection of parameters to tune was based on the arguments of the functions called in these tutorials. Methods dependant on a random seed, DCA and scVI, were run on five seeds and we averaged their metrics in order to reduce the effect of a single good or bad seed.

### UMAP plots

The UMAP plots were generated using scanpy’s implementation with default parameters, run on all the dimensions of the low dimension representation.

### Availability of data and materials

The four cell lines datasets come from [12] and were downloaded from https://github.com/LuyiTian/sc_mixology in April 2019. The TabulaMuris dataset is an in silico mixture containing all the cells from four tissues sequenced with Smart-Seq2 from the Tabula Muris consortium [22] and was downloaded with the TabulaMurisData Bioconductor package, version 1.2.0. Zhengmix4eq, Zhengmix4uneq, and Zhengmix8eq come from [11] and were downloaded from the DuoClustering2018 Bioconductor package, version 1.2.0. We generated Zhengmix5eq and Zhengmix8uneq following the same procedure in order to have more complex datasets. All datasets were subject to the same quality control pipeline, using scater [16], version 1.14.6. We removed cells three median absolute deviations (MAD) under the mean in counts and expressed genes, as well as those three MAD above the mean in percentage of mitochondrial reads.

The code and data used in this manuscript, as well as the 5, 000 pre-computed embeddings provided as a benchmark to develop new heuristics for parameter selection, are freely available under an Apache License 2.0 [30, 31].

## Competing interests

The authors declare that they have no competing interests.

## Ethics Approval

The authors declare that no ethics approval was required.

## Funding

The authors have no funding to declare.

## Authors’ contributions

All authors designed the study, analyzed the results, and wrote the manuscript. FR did all the implementation and experiments.

## Additional files

**Additional file 1**: contains additional figures and tables for the paper.

## Additional file 1

This file contains all supplementary tables and figures mentioned in the main manuscript.

**Table S1:**
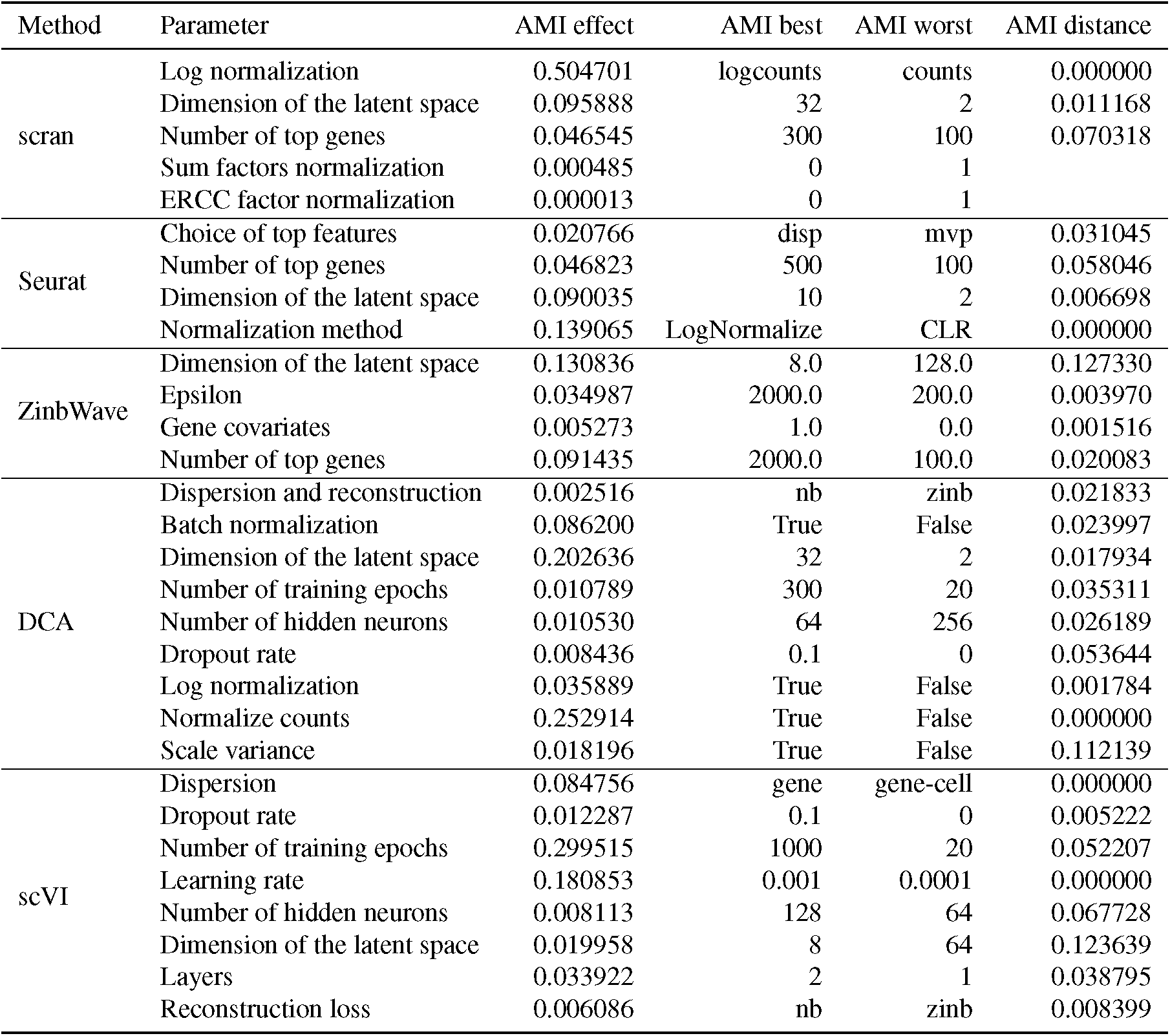
Summary of the parameter influence on the AMI. The column “AMI effect” is the maximum difference between the mean effect of the parameters on the AMI. The column “AMI best” is the parameter value with the best mean effect, and are the ones used in the “ANOVA AMI heuristic”. The column “AMI worst” is the parameter value with the worst mean effect. The column “AMI distance” is the maximum distance between the parameter values with the best effect on a dataset specific way, and the “AMI best” effect on a dataset specific way.

**Table S2:**
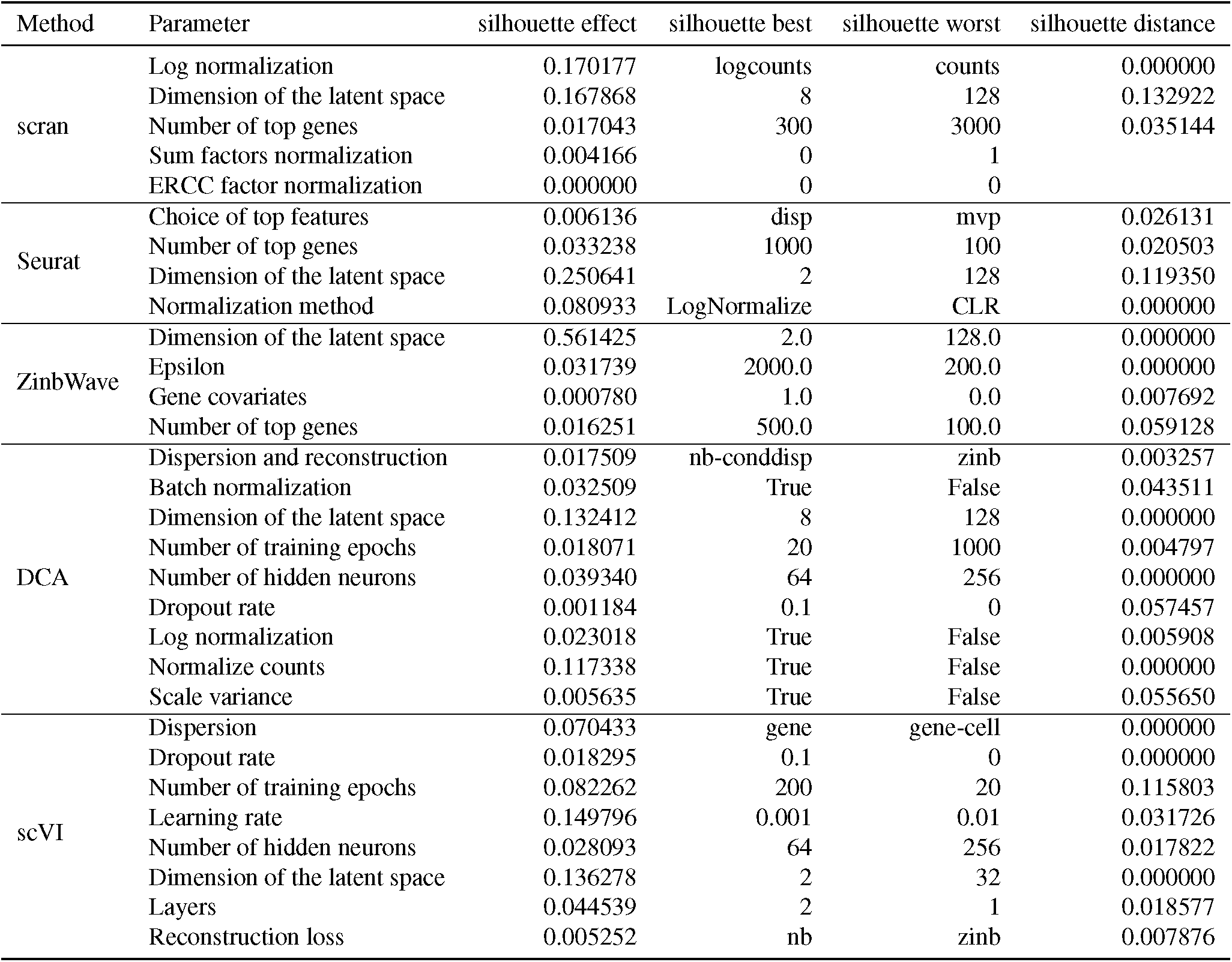
Summary of the parameter influence on the silhouette. The column “silhouette effect” is the maximum difference between the mean effect of the parameters on the silhouette. The column “silhouette best” is the parameter value with the best mean effect, and are the ones used in the “ANOVA silhouette heuristic”. The column “silhouette worst” is the parameter value with the worst mean effect. The column “silhouette distance” is the maximum distance between the parameter values with the best effect on a dataset specific way, and the “silhouette best” effect on a dataset specific way.

**Table S3:** Summary result of the ANOVA for the influence of the parameters of scran on its AMI. “SS” corresponds to the variance explained by a parameter, “df” its number of degrees of freedom, “MS” is “SS” divided by “df”, i.e. the mean variance explained by each degree of freedom, “F value”, is the observed F statistic, and “Pr(>F)” is the probability under the null hypothesis (this parameter has no influence on the AMI) to observe an F statistic this high. Each row corresponds to a factor, when they follow the format “parameter” it is the effect that this parameter has on average, when they follow the format “dataset:parameter” it is the effect of the parameter on each specific dataset (the interaction factors), “dataset” is a special one that represents the effect of the dataset on the AMI, it is used to represent the inherent complexity of the data.

|  | SS | df | MS | F value | Pr(>F) |
| --- | --- | --- | --- | --- | --- |
| scran_n_tops | 0.45 | 5.00 | 0.09 | 35.11 | 0.0000 |
| dataset | 22.93 | 9.00 | 2.55 | 982.90 | 0.0000 |
| scran_sum_factor | 0.00 | 1.00 | 0.00 | 0.05 | 0.8268 |
| scran_ercc | 0.00 | 1.00 | 0.00 | 0.00 | 0.9968 |
| scran_assay | 135.85 | 1.00 | 135.85 | 52420.33 | 0.0000 |
| scran_n_pcs | 2.06 | 7.00 | 0.29 | 113.30 | 0.0000 |
| scran_n_tops:dataset | 3.74 | 45.00 | 0.08 | 32.04 | 0.0000 |
| dataset:scran_assay | 21.32 | 9.00 | 2.37 | 913.92 | 0.0000 |
| dataset:scran_n_pcs | 0.60 | 63.00 | 0.01 | 3.70 | 0.0000 |
| Residuals | 5.60 | 2162.00 | 0.00 |  |  |

**Table S4:** Summary result of the ANOVA for the influence of the parameters of scran on its silhouette. “SS” corresponds to the variance explained by a parameter, “df” its number of degrees of freedom, “MS” is “SS” divided by “df”, i.e. the mean variance explained by each degree of freedom, “F value”, is the observed F statistic, and “Pr(>F)” is the probability under the null hypothesis (this parameter has no influence on the silhouette) to observe an F statistic this high. Each row corresponds to a factor, when they follow the format “parameter” it is the effect that this parameter has on average, when they follow the format “dataset:parameter” it is the effect of the parameter on each specific dataset (the interaction factors), “dataset” is a special one that represents the effect of the dataset on the silhouette, it is used to represent the inherent complexity of the data.

|  | SS | df | MS | F value | Pr(>F) |
| --- | --- | --- | --- | --- | --- |
| dataset | 19.50 | 9.00 | 2.17 | 591.87 | 0.0000 |
| scran_n_tops | 0.07 | 5.00 | 0.01 | 4.06 | 0.0011 |
| scran_sum_factor | 0.01 | 1.00 | 0.01 | 2.50 | 0.1138 |
| scran_ercc | 0.00 | 1.00 | 0.00 | 0.00 | 0.9974 |
| scran_assay | 15.45 | 1.00 | 15.45 | 4218.86 | 0.0000 |
| scran_n_pcs | 6.94 | 7.00 | 0.99 | 270.77 | 0.0000 |
| dataset:scran_n_tops | 0.42 | 45.00 | 0.01 | 2.57 | 0.0000 |
| dataset:scran_assay | 3.70 | 9.00 | 0.41 | 112.44 | 0.0000 |
| dataset:scran_n_pcs | 2.25 | 63.00 | 0.04 | 9.75 | 0.0000 |
| Residuals | 7.92 | 2162.00 | 0.00 |  |  |

**Table S5:** Summary result of the ANOVA for the influence of the parameters of Seurat on its AMI. “SS” corresponds to the variance explained by a parameter, “df” its number of degrees of freedom, “MS” is “SS” divided by “df”, i.e. the mean variance explained by each degree of freedom, “F value”, is the observed F statistic, and “Pr(>F)” is the probability under the null hypothesis (this parameter has no influence on the AMI) to observe an F statistic this high. Each row corresponds to a factor, when they follow the format “parameter” it is the effect that this parameter has on average, when they follow the format “dataset:parameter” it is the effect of the parameter on each specific dataset (the interaction factors), “dataset” is a special one that represents the effect of the dataset on the AMI, it is used to represent the inherent complexity of the data.

|  | SS | df | MS | F value | Pr(>F) |
| --- | --- | --- | --- | --- | --- |
| dataset | 103.26 | 9.00 | 11.47 | 2711.13 | 0.0000 |
| seurat_n_features | 0.75 | 5.00 | 0.15 | 35.53 | 0.0000 |
| seurat_n_pcs | 2.49 | 7.00 | 0.36 | 84.12 | 0.0000 |
| seurat_norm | 13.92 | 1.00 | 13.92 | 3289.15 | 0.0000 |
| seurat_find_variable | 0.23 | 2.00 | 0.12 | 27.52 | 0.0000 |
| dataset:seurat_n_features | 2.51 | 45.00 | 0.06 | 13.20 | 0.0000 |
| dataset:seurat_n_pcs | 1.12 | 63.00 | 0.02 | 4.20 | 0.0000 |
| dataset:seurat_norm | 11.95 | 9.00 | 1.33 | 313.81 | 0.0000 |
| dataset:seurat_find_variable | 1.05 | 18.00 | 0.06 | 13.84 | 0.0000 |
| Residuals | 11.51 | 2719.00 | 0.00 |  |  |

**Table S6:** Summary result of the ANOVA for the influence of the parameters of Seurat on its silhouette. “SS” corresponds to the variance explained by a parameter, “df” its number of degrees of freedom, “MS” is “SS” divided by “df”, i.e. the mean variance explained by each degree of freedom, “F value”, is the observed F statistic, and “Pr(>F)” is the probability under the null hypothesis (this parameter has no influence on the silhouette) to observe an F statistic this high. Each row corresponds to a factor, when they follow the format “parameter” it is the effect that this parameter has on average, when they follow the format “dataset:parameter” it is the effect of the parameter on each specific dataset (the interaction factors), “dataset” is a special one that represents the effect of the dataset on the silhouette, it is used to represent the inherent complexity of the data.

|  | SS | df | MS | F value | Pr(>F) |
| --- | --- | --- | --- | --- | --- |
| dataset | 62.00 | 9.00 | 6.89 | 2855.27 | 0.0000 |
| seurat_n_features | 0.36 | 5.00 | 0.07 | 30.15 | 0.0000 |
| seurat_n_pcs | 21.87 | 7.00 | 3.12 | 1295.01 | 0.0000 |
| seurat_norm | 4.71 | 1.00 | 4.71 | 1953.86 | 0.0000 |
| seurat_find_variable | 0.02 | 2.00 | 0.01 | 3.92 | 0.0200 |
| dataset:seurat_n_features | 0.51 | 45.00 | 0.01 | 4.66 | 0.0000 |
| dataset:seurat_n_pcs | 7.03 | 63.00 | 0.11 | 46.28 | 0.0000 |
| dataset:seurat_norm | 0.57 | 9.00 | 0.06 | 26.12 | 0.0000 |
| dataset:seurat_find_variable | 0.42 | 18.00 | 0.02 | 9.56 | 0.0000 |
| Residuals | 6.56 | 2719.00 | 0.00 |  |  |

**Table S7:** Summary result of the ANOVA for the influence of the parameters of ZinbWave on its AMI. “SS” corresponds to the variance explained by a parameter, “df” its number of degrees of freedom, “MS” is “SS” divided by “df”, i.e. the mean variance explained by each degree of freedom, “F value”, is the observed F statistic, and “Pr(>F)” is the probability under the null hypothesis (this parameter has no influence on the AMI) to observe an F statistic this high. Each row corresponds to a factor, when they follow the format “parameter” it is the effect that this parameter has on average, when they follow the format “dataset:parameter” it is the effect of the parameter on each specific dataset (the interaction factors), “dataset” is a special one that represents the effect of the dataset on the AMI, it is used to represent the inherent complexity of the data.

|  | SS | df | MS | F value | Pr(>F) |
| --- | --- | --- | --- | --- | --- |
| dataset | 186.02 | 9.00 | 20.67 | 6293.87 | 0.0000 |
| zinbwave_dims | 5.10 | 7.00 | 0.73 | 221.82 | 0.0000 |
| zinbwave_epsilon | 0.51 | 3.00 | 0.17 | 51.78 | 0.0000 |
| zinbwave_gene_covariate | 0.02 | 1.00 | 0.02 | 6.49 | 0.0109 |
| zinbwave_keep_variance | 2.65 | 4.00 | 0.66 | 201.45 | 0.0000 |
| dataset:zinbwave_dims | 12.67 | 63.00 | 0.20 | 61.22 | 0.0000 |
| dataset:zinbwave_epsilon | 0.81 | 27.00 | 0.03 | 9.14 | 0.0000 |
| dataset:zinbwave_gene_covariate | 0.03 | 9.00 | 0.00 | 0.93 | 0.5013 |
| dataset:zinbwave_keep_variance | 2.18 | 36.00 | 0.06 | 18.45 | 0.0000 |
| Residuals | 9.56 | 2910.00 | 0.00 |  |  |

**Table S8:**
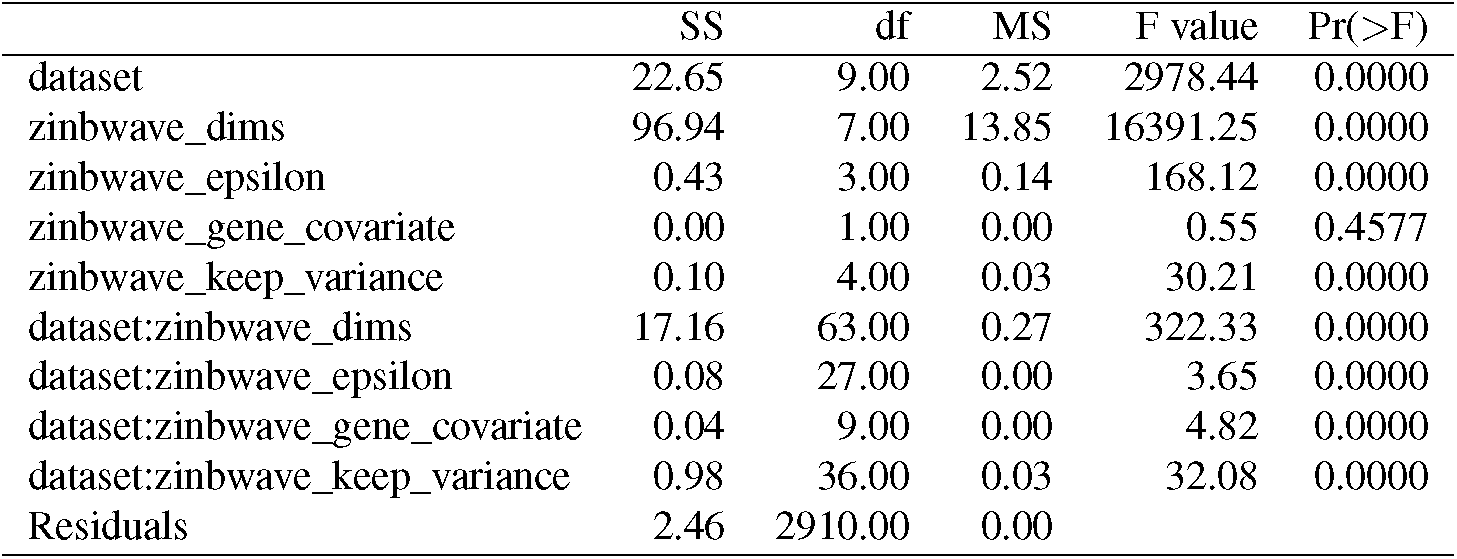
Summary result of the ANOVA for the influence of the parameters of ZinbWave on its silhouette. “SS” corresponds to the variance explained by a parameter, “df” its number of degrees of freedom, “MS” is “SS” divided by “df”, i.e. the mean variance explained by each degree of freedom, “F value”, is the observed F statistic, and “Pr(>F)” is the probability under the null hypothesis (this parameter has no influence on the silhouette) to observe an F statistic this high. Each row corresponds to a factor, when they follow the format “parameter” it is the effect that this parameter has on average, when they follow the format “dataset:parameter” it is the effect of the parameter on each specific dataset (the interaction factors), “dataset” is a special one that represents the effect of the dataset on the silhouette, it is used to represent the inherent complexity of the data.

**Table S9:**
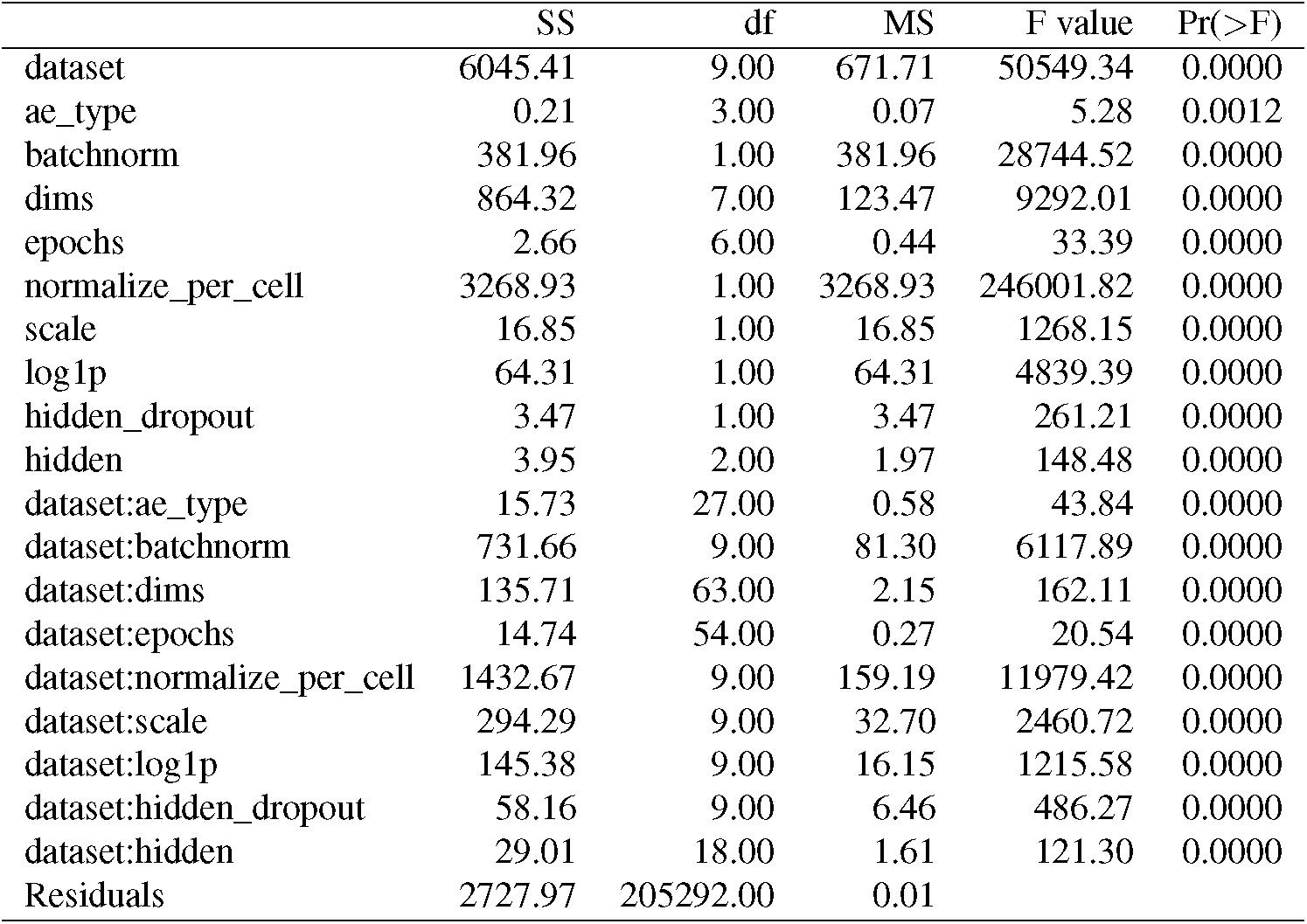
Summary result of the ANOVA for the influence of the parameters of DCA on its AMI. “SS” corresponds to the variance explained by a parameter, “df” its number of degrees of freedom, “MS” is “SS” divided by “df”, i.e. the mean variance explained by each degree of freedom, “F value”, is the observed F statistic, and “Pr(>F)” is the probability under the null hypothesis (this parameter has no influence on the AMI) to observe an F statistic this high. Each row corresponds to a factor, when they follow the format “parameter” it is the effect that this parameter has on average, when they follow the format “dataset:parameter” it is the effect of the parameter on each specific dataset (the interaction factors), “dataset” is a special one that represents the effect of the dataset on the AMI, it is used to represent the inherent complexity of the data.

|  | SS | df | MS | F value | Pr(>F) |
| --- | --- | --- | --- | --- | --- |
| dataset | 6045.41 | 9.00 | 671.71 | 50549.34 | 0.0000 |
| ae_type | 0.21 | 3.00 | 0.07 | 5.28 | 0.0012 |
| batchnorm | 381.96 | 1.00 | 381.96 | 28744.52 | 0.0000 |
| dims | 864.32 | 7.00 | 123.47 | 9292.01 | 0.0000 |
| epochs | 2.66 | 6.00 | 0.44 | 33.39 | 0.0000 |
| normalize_per_cell | 3268.93 | 1.00 | 3268.93 | 246001.82 | 0.0000 |
| scale | 16.85 | 1.00 | 16.85 | 1268.15 | 0.0000 |
| log1p | 64.31 | 1.00 | 64.31 | 4839.39 | 0.0000 |
| hidden_dropout | 3.47 | 1.00 | 3.47 | 261.21 | 0.0000 |
| hidden | 3.95 | 2.00 | 1.97 | 148.48 | 0.0000 |
| dataset:ae_type | 15.73 | 27.00 | 0.58 | 43.84 | 0.0000 |
| dataset:batchnorm | 731.66 | 9.00 | 81.30 | 6117.89 | 0.0000 |
| dataset:dims | 135.71 | 63.00 | 2.15 | 162.11 | 0.0000 |
| dataset:epochs | 14.74 | 54.00 | 0.27 | 20.54 | 0.0000 |
| dataset:normalize_per_cell | 1432.67 | 9.00 | 159.19 | 11979.42 | 0.0000 |
| dataset:scale | 294.29 | 9.00 | 32.70 | 2460.72 | 0.0000 |
| dataset:log1p | 145.38 | 9.00 | 16.15 | 1215.58 | 0.0000 |
| dataset:hidden_dropout | 58.16 | 9.00 | 6.46 | 486.27 | 0.0000 |
| dataset:hidden | 29.01 | 18.00 | 1.61 | 121.30 | 0.0000 |
| Residuals | 2727.97 | 205292.00 | 0.01 |  |  |

**Table S10:** Summary result of the ANOVA for the influence of the parameters of DCA on its silhouette. “SS” corresponds to the variance explained by a parameter, “df” its number of degrees of freedom, “MS” is “SS” divided by “df”, i.e. the mean variance explained by each degree of freedom, “F value”, is the observed F statistic, and “Pr(>F)” is the probability under the null hypothesis (this parameter has no influence on the silhouette) to observe an F statistic this high. Each row corresponds to a factor, when they follow the format “parameter” it is the effect that this parameter has on average, when they follow the format “dataset:parameter” it is the effect of the parameter on each specific dataset (the interaction factors), “dataset” is a special one that represents the effect of the dataset on the silhouette, it is used to represent the inherent complexity of the data.

|  | SS | df | MS | F value | Pr(>F) |
| --- | --- | --- | --- | --- | --- |
| dataset | 3542.22 | 9.00 | 393.58 | 46229.81 | 0.0000 |
| ae_type | 9.36 | 3.00 | 3.12 | 366.43 | 0.0000 |
| batchnorm | 53.66 | 1.00 | 53.66 | 6303.34 | 0.0000 |
| dims | 396.37 | 7.00 | 56.62 | 6651.01 | 0.0000 |
| epochs | 7.71 | 6.00 | 1.28 | 150.92 | 0.0000 |
| normalize_per_cell | 700.98 | 1.00 | 700.98 | 82336.34 | 0.0000 |
| scale | 1.56 | 1.00 | 1.56 | 183.13 | 0.0000 |
| log1p | 26.88 | 1.00 | 26.88 | 3157.50 | 0.0000 |
| hidden_dropout | 0.06 | 1.00 | 0.06 | 7.41 | 0.0065 |
| hidden | 52.00 | 2.00 | 26.00 | 3054.02 | 0.0000 |
| dataset:ae_type | 7.71 | 27.00 | 0.29 | 33.56 | 0.0000 |
| dataset:batchnorm | 494.51 | 9.00 | 54.95 | 6453.93 | 0.0000 |
| dataset:dims | 124.46 | 63.00 | 1.98 | 232.05 | 0.0000 |
| dataset:epochs | 8.94 | 54.00 | 0.17 | 19.46 | 0.0000 |
| dataset:normalize_per_cell | 131.01 | 9.00 | 14.56 | 1709.83 | 0.0000 |
| dataset:scale | 142.09 | 9.00 | 15.79 | 1854.47 | 0.0000 |
| dataset:log1p | 72.51 | 9.00 | 8.06 | 946.32 | 0.0000 |
| dataset:hidden_dropout | 37.63 | 9.00 | 4.18 | 491.17 | 0.0000 |
| dataset:hidden | 14.03 | 18.00 | 0.78 | 91.53 | 0.0000 |
| Residuals | 1747.77 | 205292.00 | 0.01 |  |  |

**Table S11:** Summary result of the ANOVA for the influence of the parameters of scVI on its AMI. “SS” corresponds to the variance explained by a parameter, “df” its number of degrees of freedom, “MS” is “SS” divided by “df”, i.e. the mean variance explained by each degree of freedom, “F value”, is the observed F statistic, and “Pr(>F)” is the probability under the null hypothesis (this parameter has no influence on the AMI) to observe an F statistic this high. Each row corresponds to a factor, when they follow the format “parameter” it is the effect that this parameter has on average, when they follow the format “dataset:parameter” it is the effect of the parameter on each specific dataset (the interaction factors), “dataset” is a special one that represents the effect of the dataset on the AMI, it is used to represent the inherent complexity of the data.

|  | SS | df | MS | F value | Pr(>F) |
| --- | --- | --- | --- | --- | --- |
| n_latent | 4.07 | 7.00 | 0.58 | 12.56 | 0.0000 |
| dataset | 3304.57 | 9.00 | 367.17 | 7934.53 | 0.0000 |
| epochs | 836.88 | 6.00 | 139.48 | 3014.13 | 0.0000 |
| dispersion | 143.17 | 1.00 | 143.17 | 3093.80 | 0.0000 |
| n_layers | 23.07 | 1.00 | 23.07 | 498.51 | 0.0000 |
| n_hidden | 0.90 | 2.00 | 0.45 | 9.78 | 0.0001 |
| dropout_rate | 2.98 | 1.00 | 2.98 | 64.44 | 0.0000 |
| lr | 471.85 | 2.00 | 235.92 | 5098.25 | 0.0000 |
| reconstruction_loss | 0.76 | 1.00 | 0.76 | 16.35 | 0.0001 |
| n_latent:dataset | 39.76 | 63.00 | 0.63 | 13.64 | 0.0000 |
| dataset:epochs | 383.74 | 54.00 | 7.11 | 153.57 | 0.0000 |
| dataset:dispersion | 240.35 | 9.00 | 26.71 | 577.09 | 0.0000 |
| dataset:n_layers | 54.17 | 9.00 | 6.02 | 130.08 | 0.0000 |
| dataset:n_hidden | 61.33 | 18.00 | 3.41 | 73.63 | 0.0000 |
| dataset:dropout_rate | 5.08 | 9.00 | 0.56 | 12.20 | 0.0000 |
| dataset:lr | 657.45 | 18.00 | 36.53 | 789.30 | 0.0000 |
| dataset:reconstruction_loss | 2.88 | 9.00 | 0.32 | 6.91 | 0.0000 |
| Residuals | 3679.78 | 79519.00 | 0.05 |  |  |

**Table S12:** Summary result of the ANOVA for the influence of the parameters of scVI on its silhouette. “SS” corresponds to the variance explained by a parameter, “df” its number of degrees of freedom, “MS” is “SS” divided by “df”, i.e. the mean variance explained by each degree of freedom, “F value”, is the observed F statistic, and “Pr(>F)” is the probability under the null hypothesis (this parameter has no influence on the silhouette) to observe an F statistic this high. Each row corresponds to a factor, when they follow the format “parameter” it is the effect that this parameter has on average, when they follow the format “dataset:parameter” it is the effect of the parameter on each specific dataset (the interaction factors), “dataset” is a special one that represents the effect of the dataset on the silhouette, it is used to represent the inherent complexity of the data.

|  | SS | df | MS | F value | Pr(>F) |
| --- | --- | --- | --- | --- | --- |
| n_latent | 151.33 | 7.00 | 21.62 | 618.87 | 0.0000 |
| dataset | 914.48 | 9.00 | 101.61 | 2908.80 | 0.0000 |
| epochs | 61.21 | 6.00 | 10.20 | 292.04 | 0.0000 |
| dispersion | 98.93 | 1.00 | 98.93 | 2832.20 | 0.0000 |
| n_layers | 39.34 | 1.00 | 39.34 | 1126.30 | 0.0000 |
| n_hidden | 10.57 | 2.00 | 5.28 | 151.29 | 0.0000 |
| dropout_rate | 6.65 | 1.00 | 6.65 | 190.50 | 0.0000 |
| lr | 302.15 | 2.00 | 151.07 | 4324.90 | 0.0000 |
| reconstruction_loss | 0.53 | 1.00 | 0.53 | 15.27 | 0.0001 |
| n_latent:dataset | 56.63 | 63.00 | 0.90 | 25.73 | 0.0000 |
| dataset:epochs | 293.43 | 54.00 | 5.43 | 155.56 | 0.0000 |
| dataset:dispersion | 130.38 | 9.00 | 14.49 | 414.73 | 0.0000 |
| dataset:n_layers | 27.01 | 9.00 | 3.00 | 85.91 | 0.0000 |
| dataset:n_hidden | 32.82 | 18.00 | 1.82 | 52.19 | 0.0000 |
| dataset:dropout_rate | 1.94 | 9.00 | 0.22 | 6.18 | 0.0000 |
| dataset:lr | 410.50 | 18.00 | 22.81 | 652.87 | 0.0000 |
| dataset:reconstruction_loss | 0.91 | 9.00 | 0.10 | 2.88 | 0.0021 |
| Residuals | 2777.70 | 79519.00 | 0.03 |  |  |

**Figure S1:**
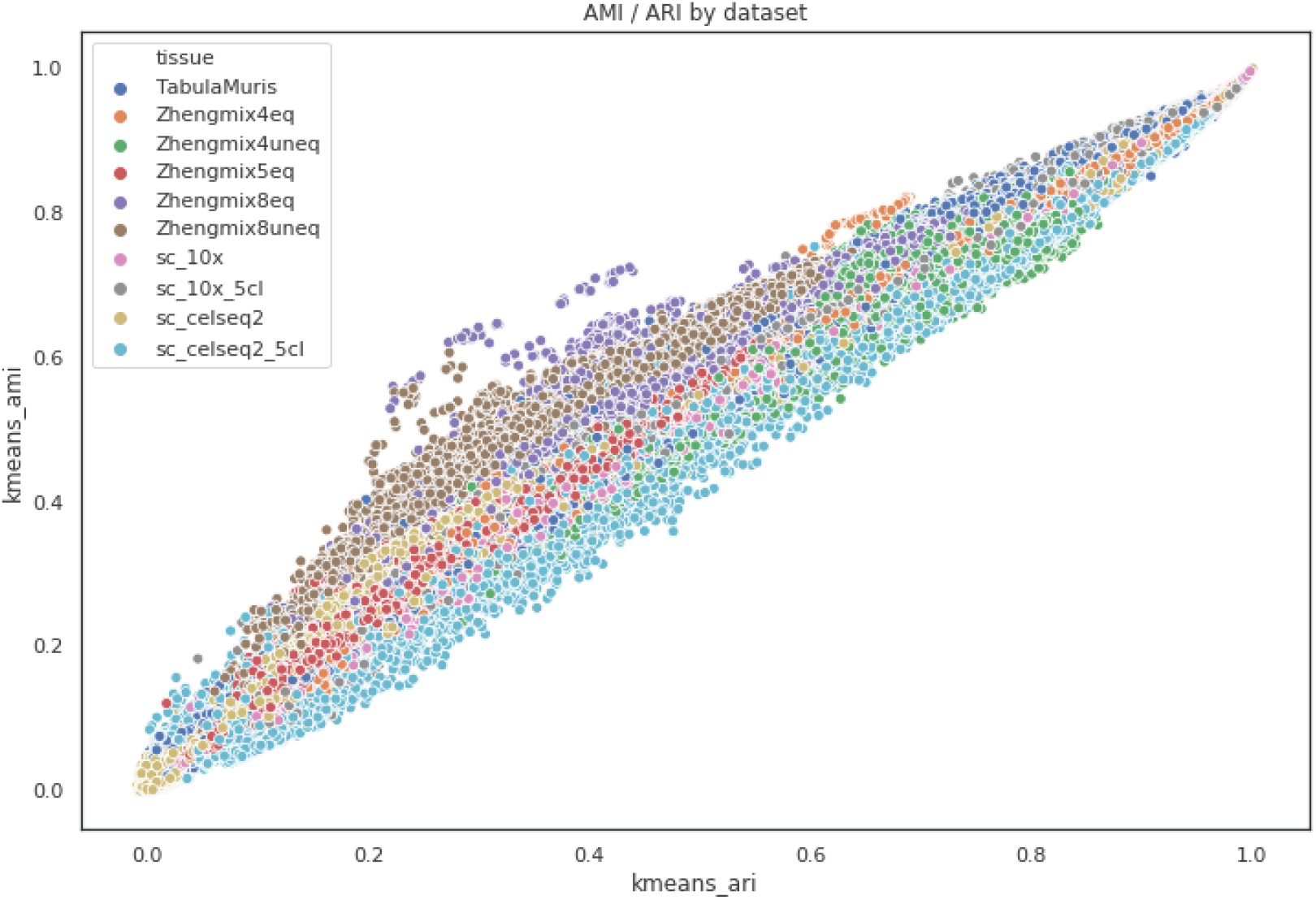
Relationship between AMI (vertical axis) and ARI (horizontal axis), computed with *k*-means clustering, colored by dataset. Each point represents the result of one experiment (running one method with a particular set of parameters on one dataset). We see that, overall, ARI and AMI are strongly correlated, particularly for a given dataset.

**Figure S2:**
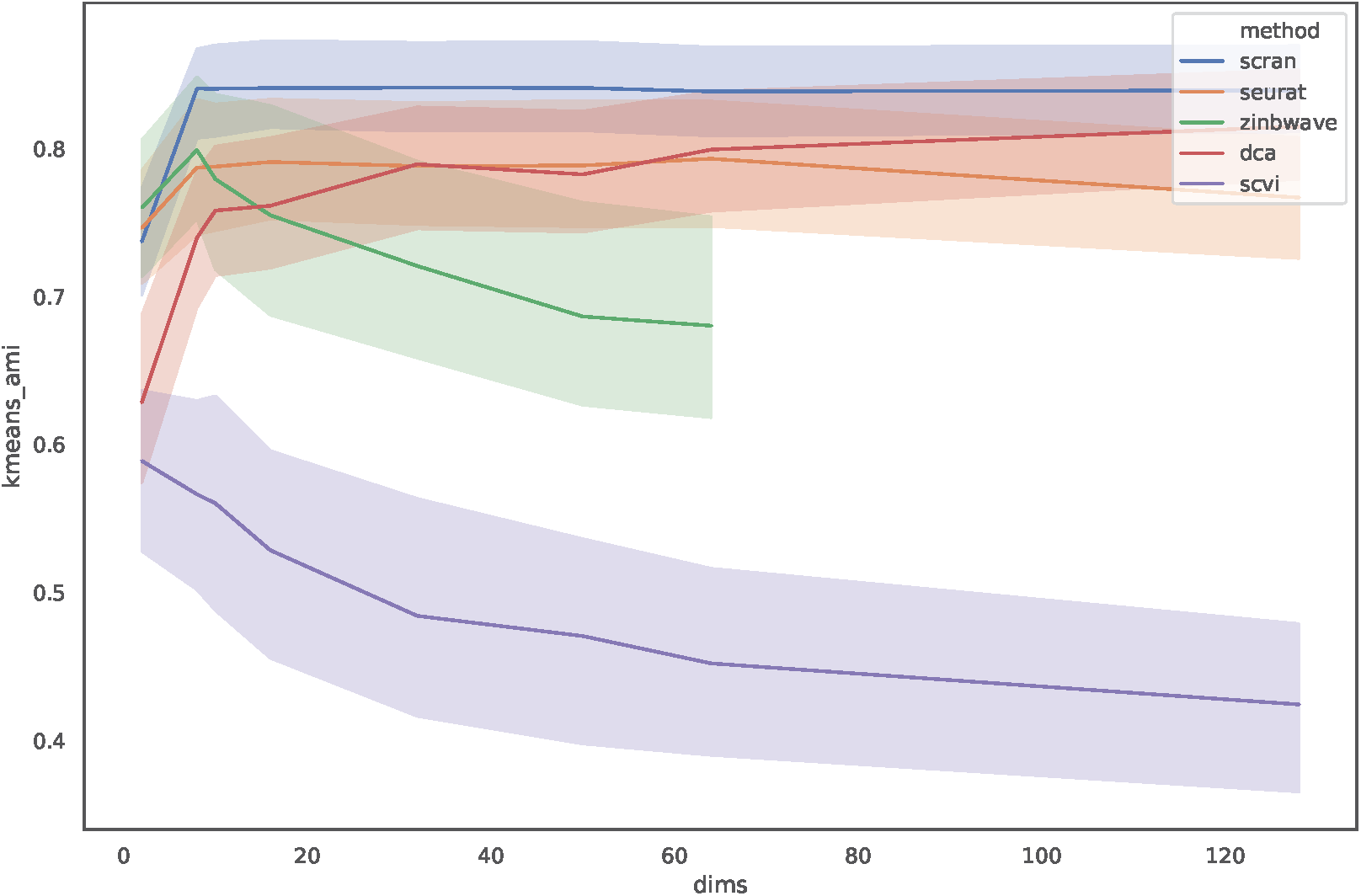
Mean AMI score across the 10 datasets (solid lines) for each method with default parameters. The transparent lines are the 95% confidence interval, which is large since we only have samples per point. The x axis is the dimension of the latent space, in order to observe its effect on the AMI.

**Figure S3:**
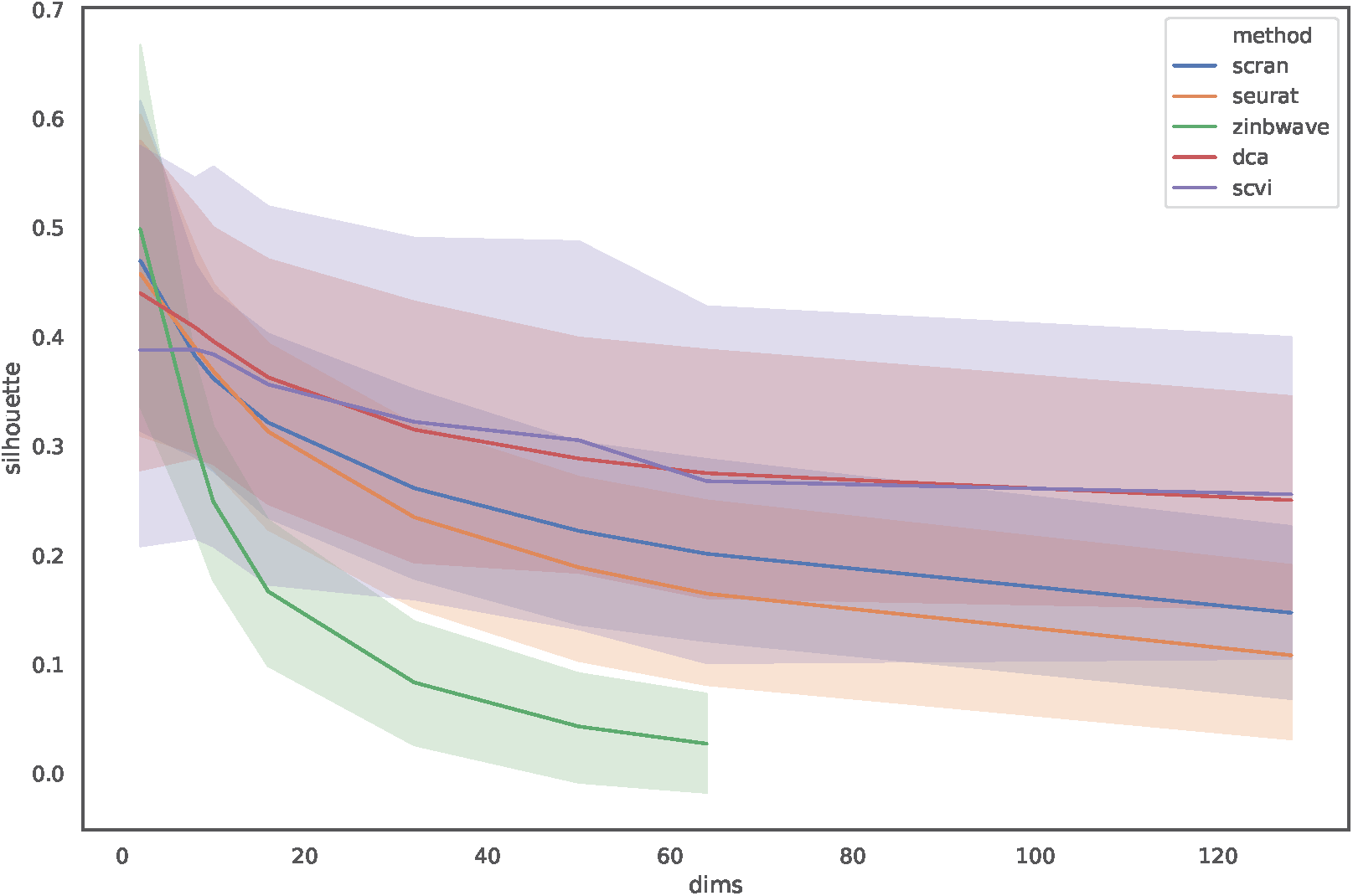
Mean silhouette score across the 10 datasets (solid lines) for each method with default parameters. The transparent lines are the 95% confidence interval, which is large since we only have samples per point. The x axis is the dimension of the latent space, in order to observe its effect on the silhouette.

**Figure S4:**
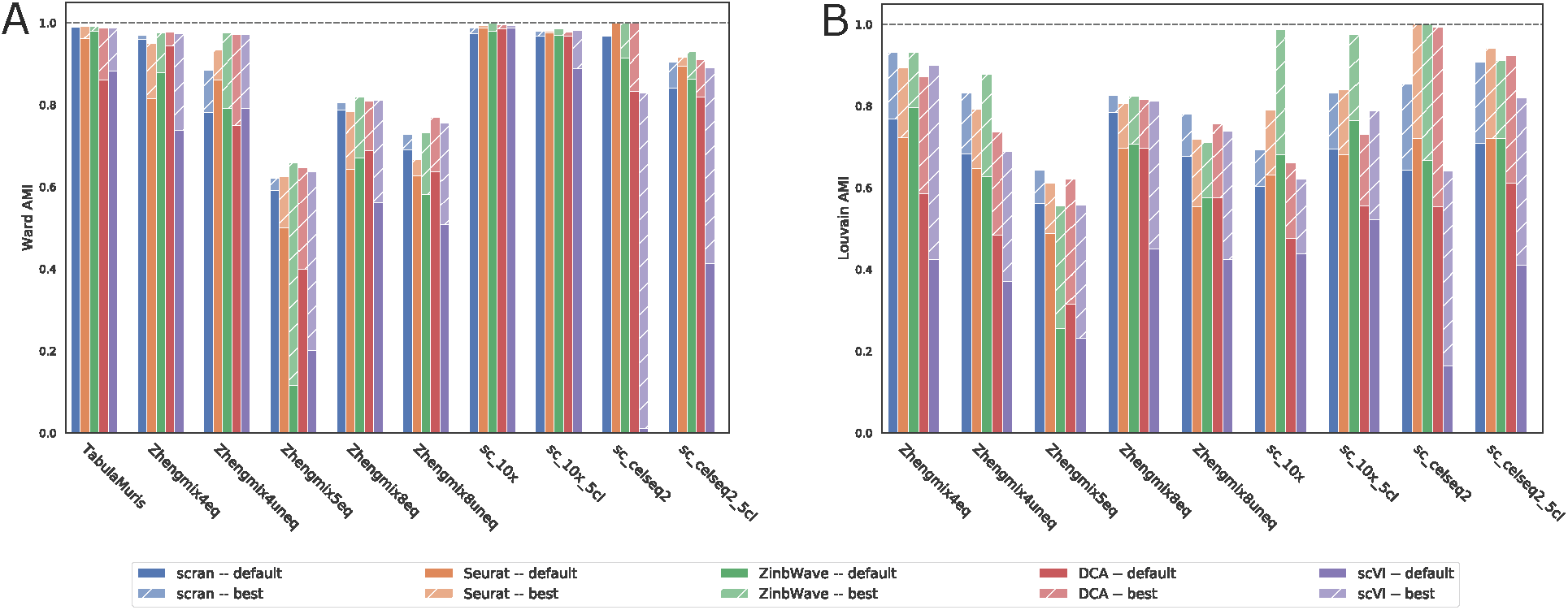
AMI after **A**. Ward clustering and **B**. Louvain clustering (right) of five DR pipelines (scran, Seurat, ZinbWave, DCA and scVI) with default parameters and a dimension of 10 (legend “default”) or after parameter optimization (legend “best”) on our benchmark of ten datasets.

**Figure S5:**
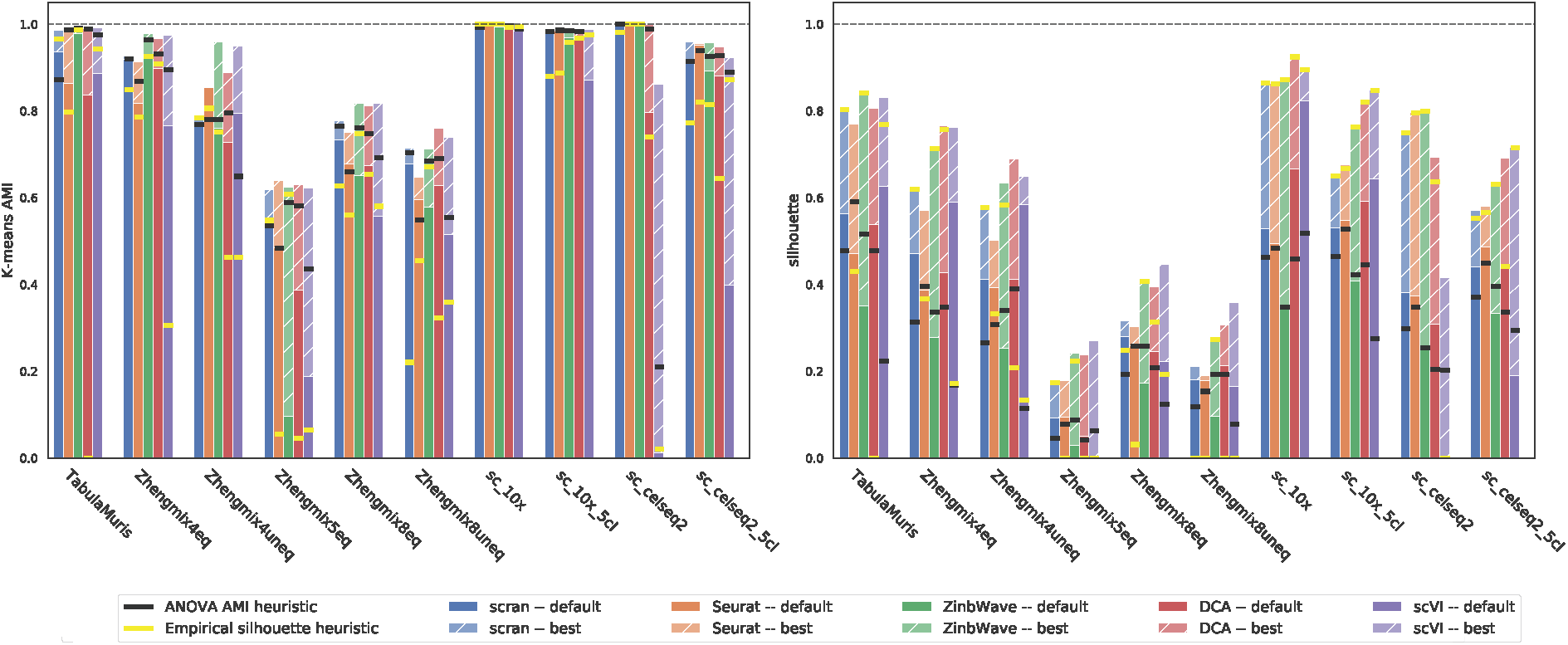
Performance of five DR pipelines (scran, Seurat, ZinbWave, DCA and scVI) with default parameters and a dimension of 10 (legend “default”) or after parameter optimization (legend “best”) on our benchmark of ten dataset. The “ANOVA AMI heuristic” corresponds to the performances of the new default parameters. The “Empirical silhouette heuristic” corresponds to the performance of the heuristic using the best empirical silhouette.

**Figure S6:**
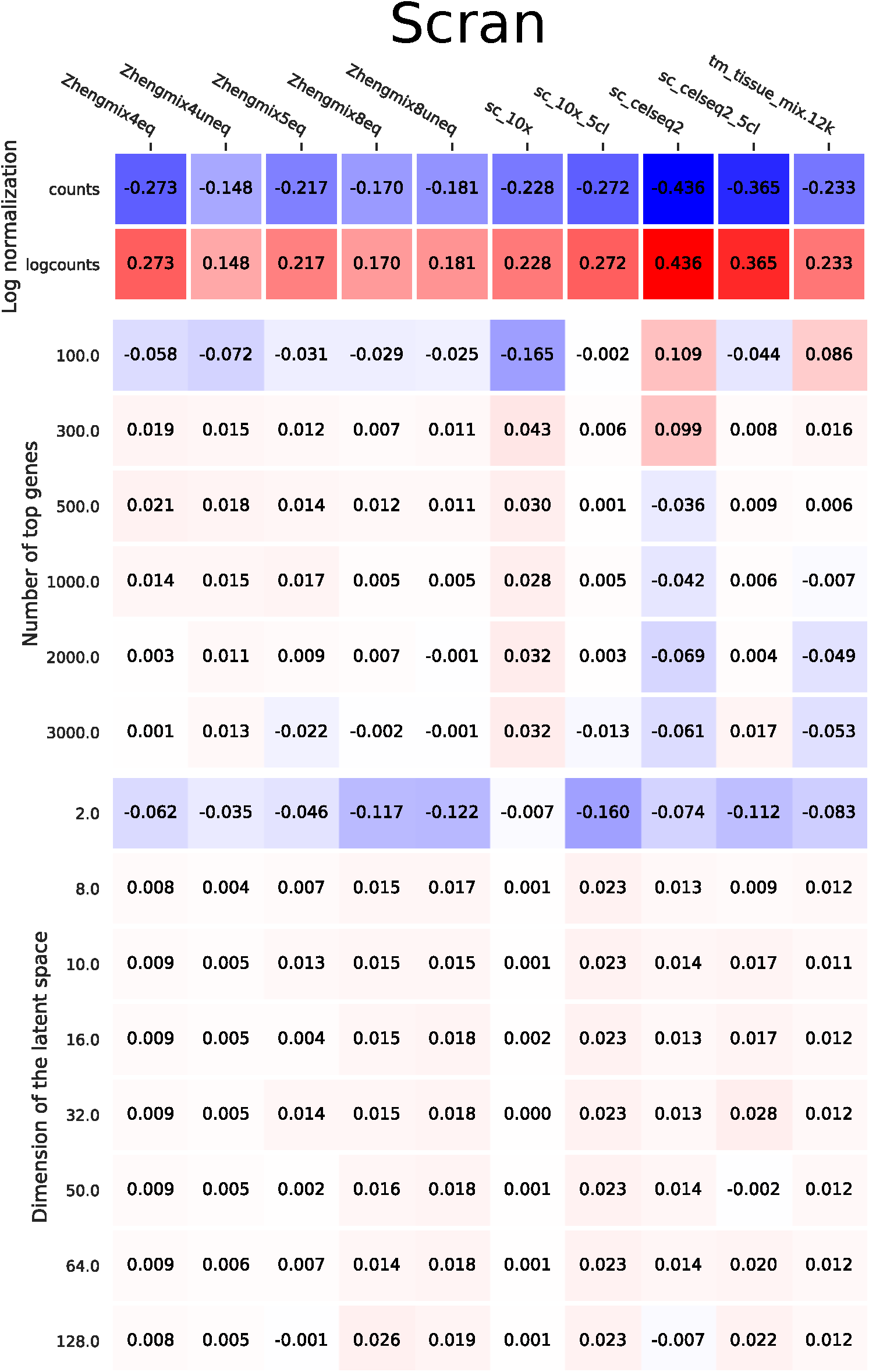
Heatmap visualization of the mean effect of each parameter value of scran on its AMI. Each columns corresponds to a dataset. The rows are split by parameter and their values, the numbers show the average effect of that parameter value on the AMI compared to the mean AMI for scran on that dataset. These effects come from a factorial ANOVA.

**Figure S7:**
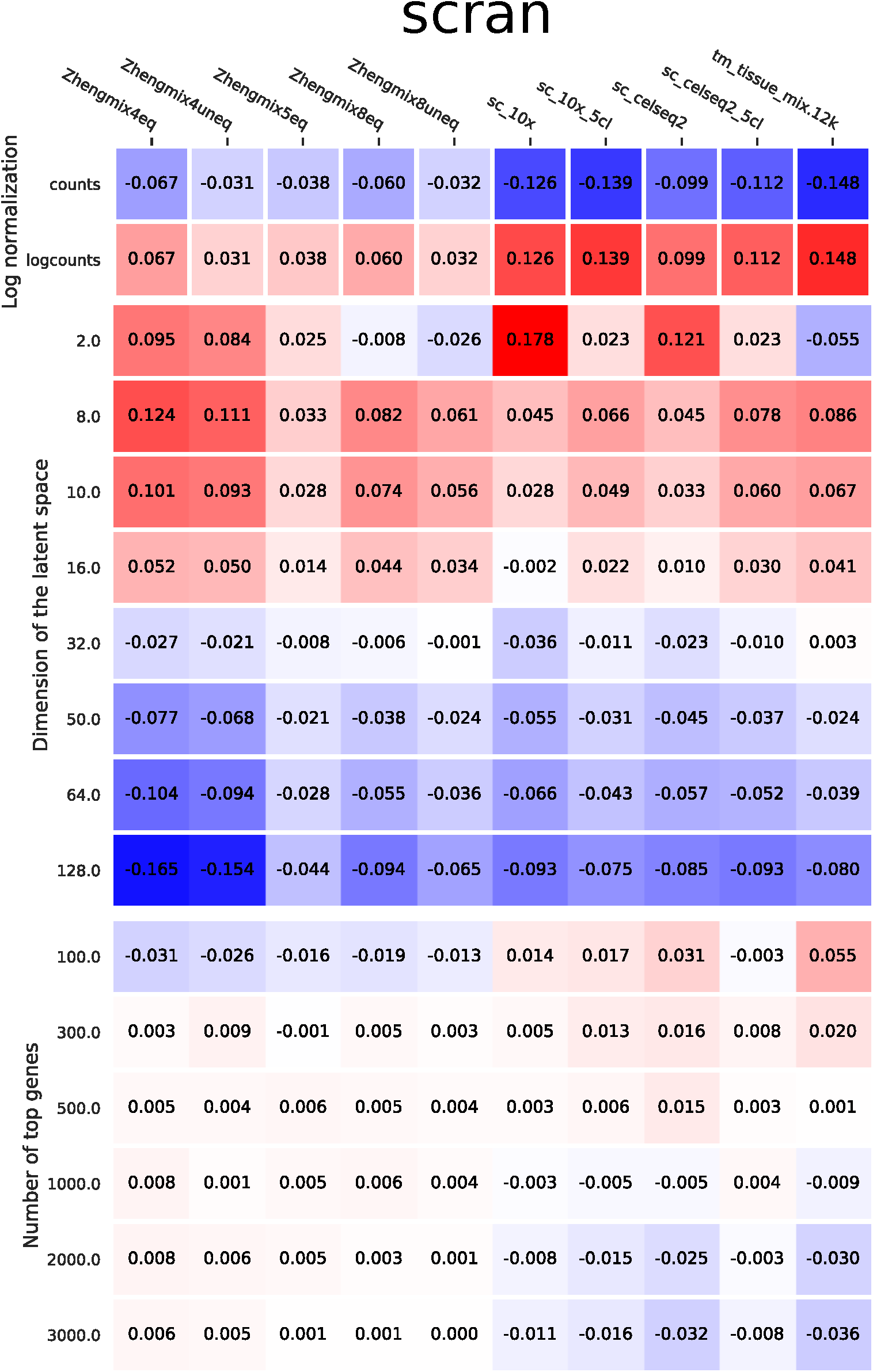
Heatmap visualization of the mean effect of each parameter value of scran on its silhouette. Each columns corresponds to a dataset. The rows are split by parameter and their values, the numbers show the average effect of that parameter value on the silhouette compared to the mean silhouette for scran on that dataset. These effects come from a factorial ANOVA.

**Figure S8:**
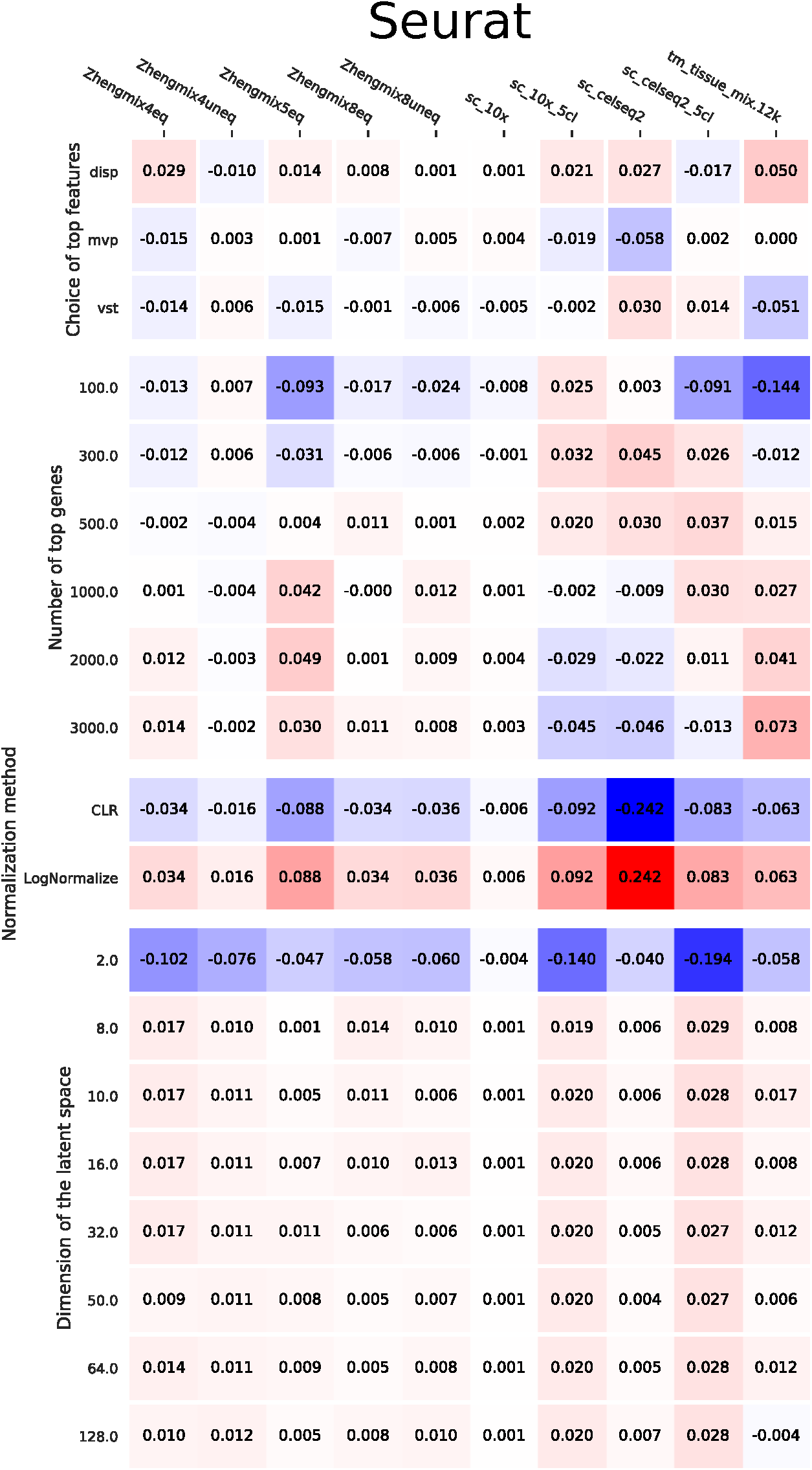
Heatmap visualization of the mean effect of each parameter value of Seurat on its AMI. Each columns corresponds to a dataset. The rows are split by parameter and their values, the numbers show the average effect of that parameter value on the AMI compared to the mean AMI for Seurat on that dataset. These effects come from a factorial ANOVA.

**Figure S9:**
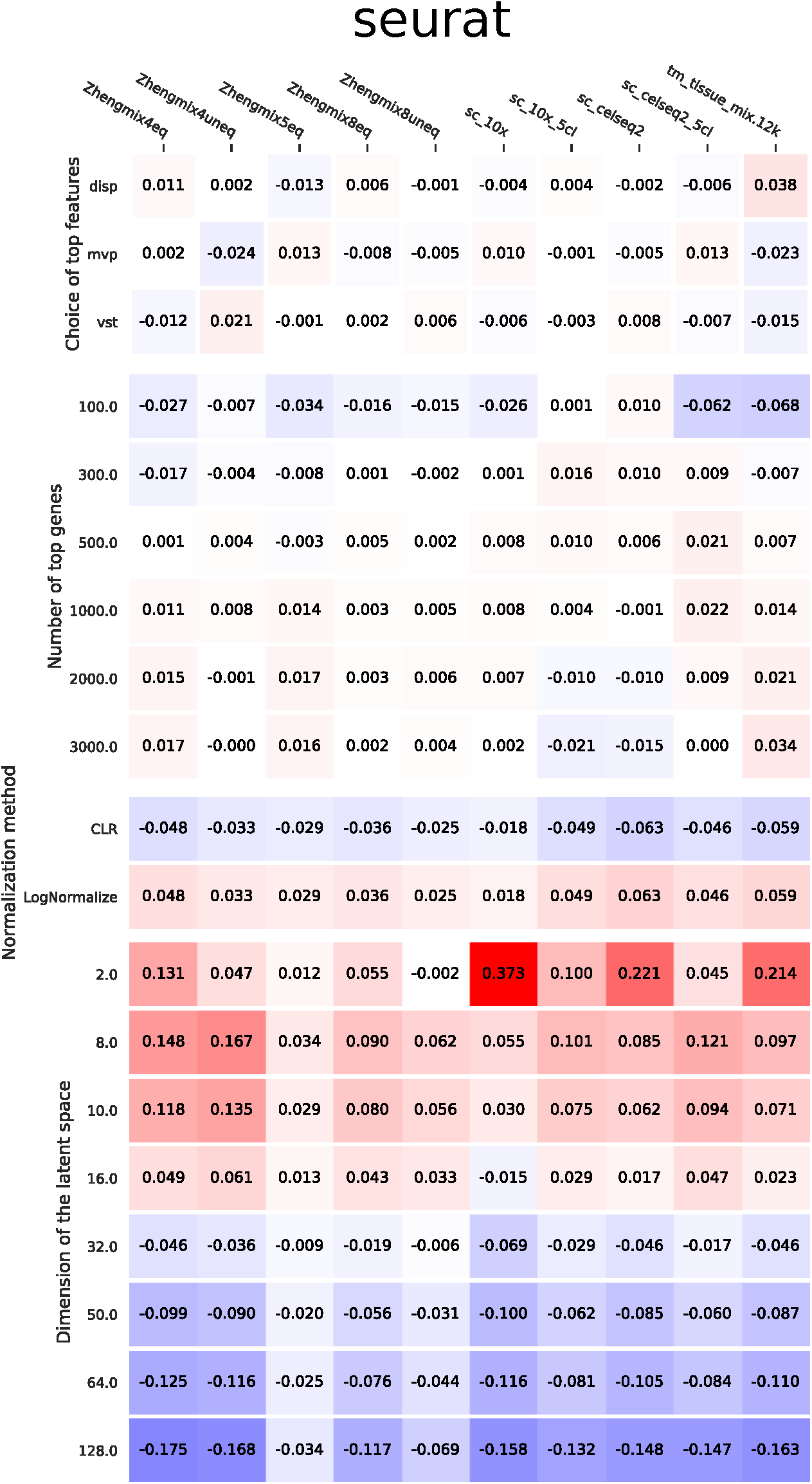
Heatmap visualization of the mean effect of each parameter value of Seurat on its silhouette. Each columns corresponds to a dataset. The rows are split by parameter and their values, the numbers show the average effect of that parameter value on the silhouette compared to the mean silhouette for Seurat on that dataset. These effects come from a factorial ANOVA.

**Figure S10:**
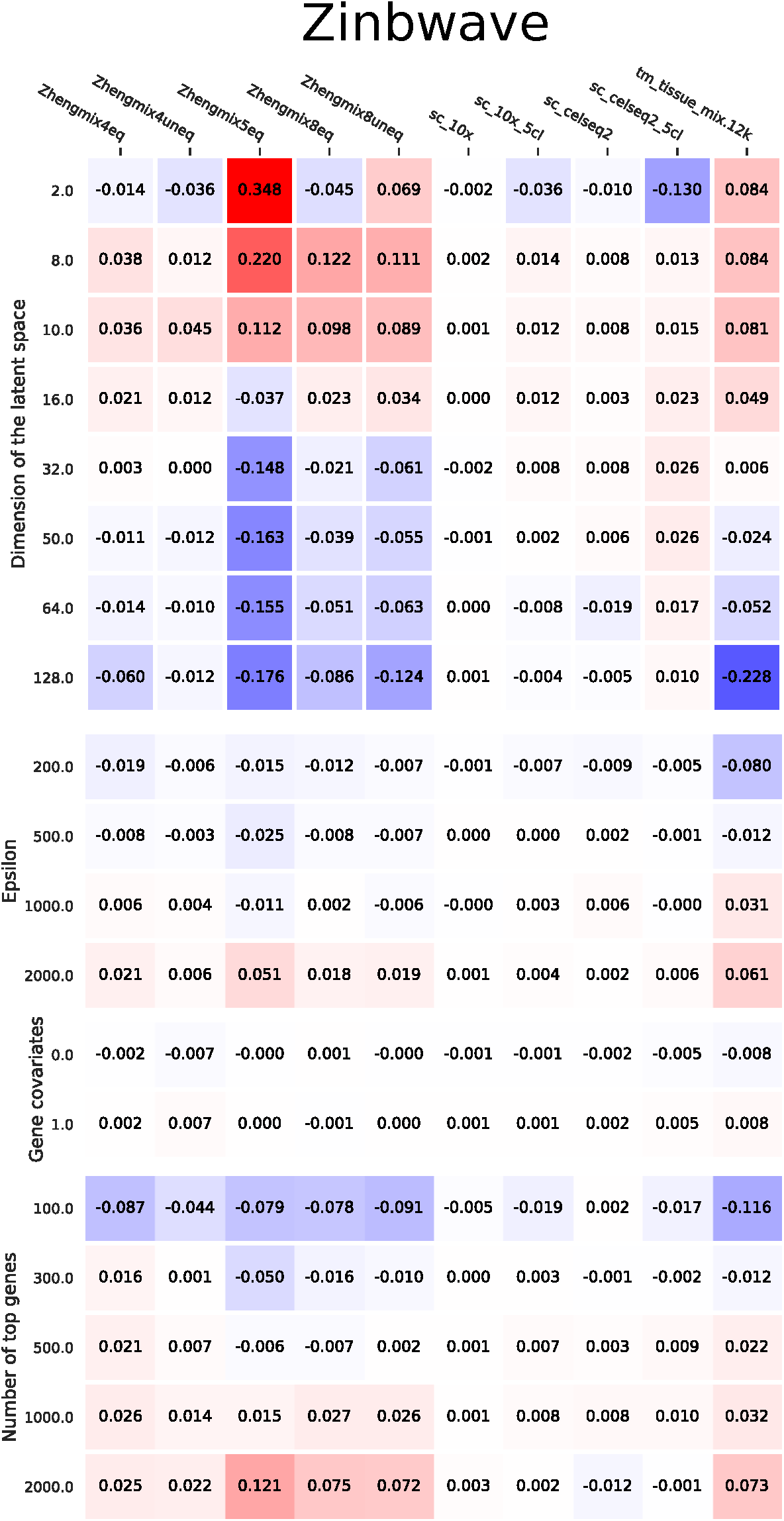
Heatmap visualization of the mean effect of each parameter value of ZinbWave on its AMI. Each columns corresponds to a dataset. The rows are split by parameter and their values, the numbers show the average effect of that parameter value on the AMI compared to the mean AMI for ZinbWave on that dataset. These effects come from a factorial ANOVA.

**Figure S11:**
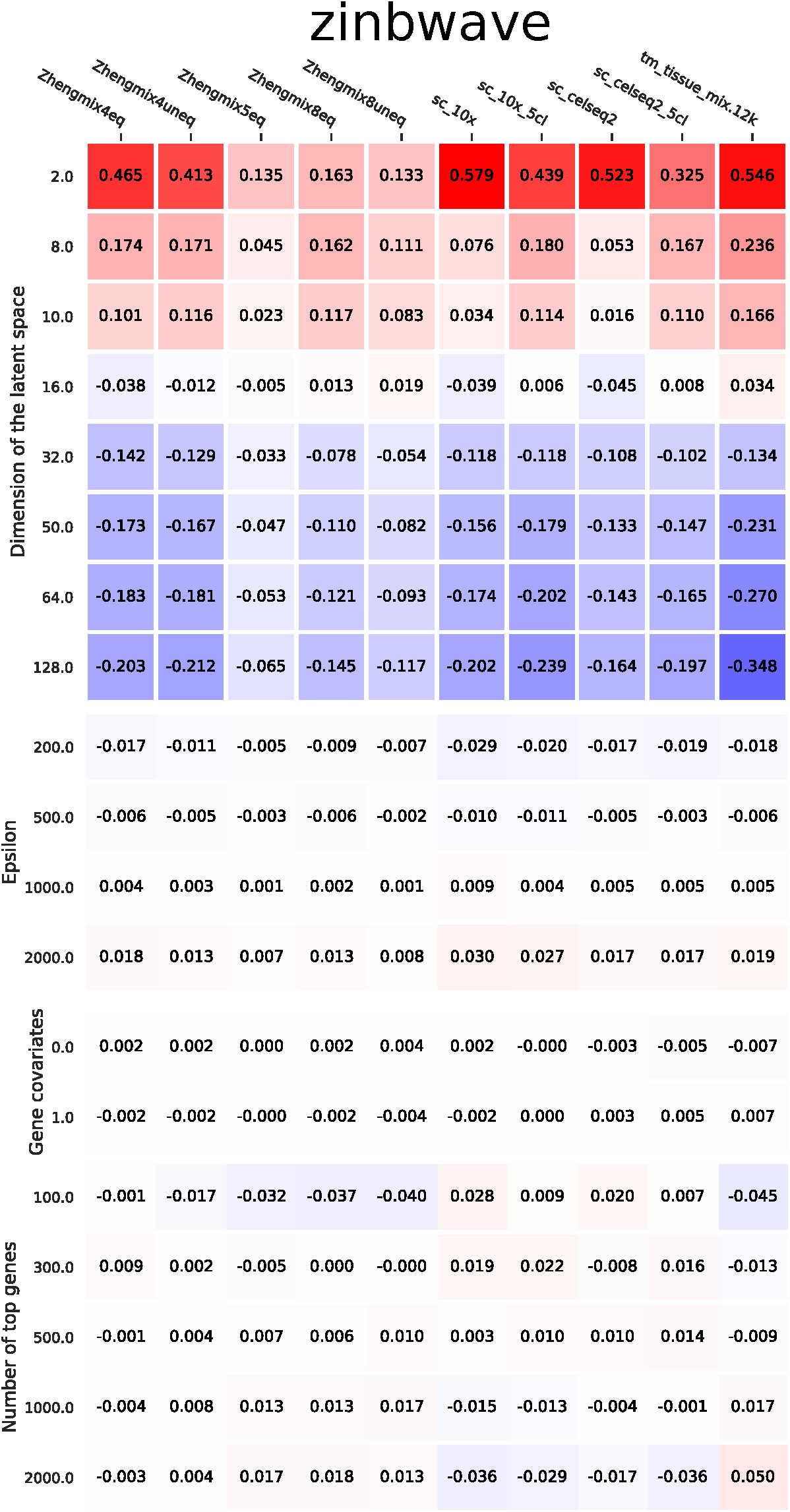
Heatmap visualization of the mean effect of each parameter value of ZinbWave on its silhouette Each columns corresponds to a dataset. The rows are split by parameter and their values, the numbers show the average effect of that parameter value on the silhouette compared to the mean silhouette for ZinbWave on that dataset. These effects come from a factorial ANOVA.

**Figure S12:**
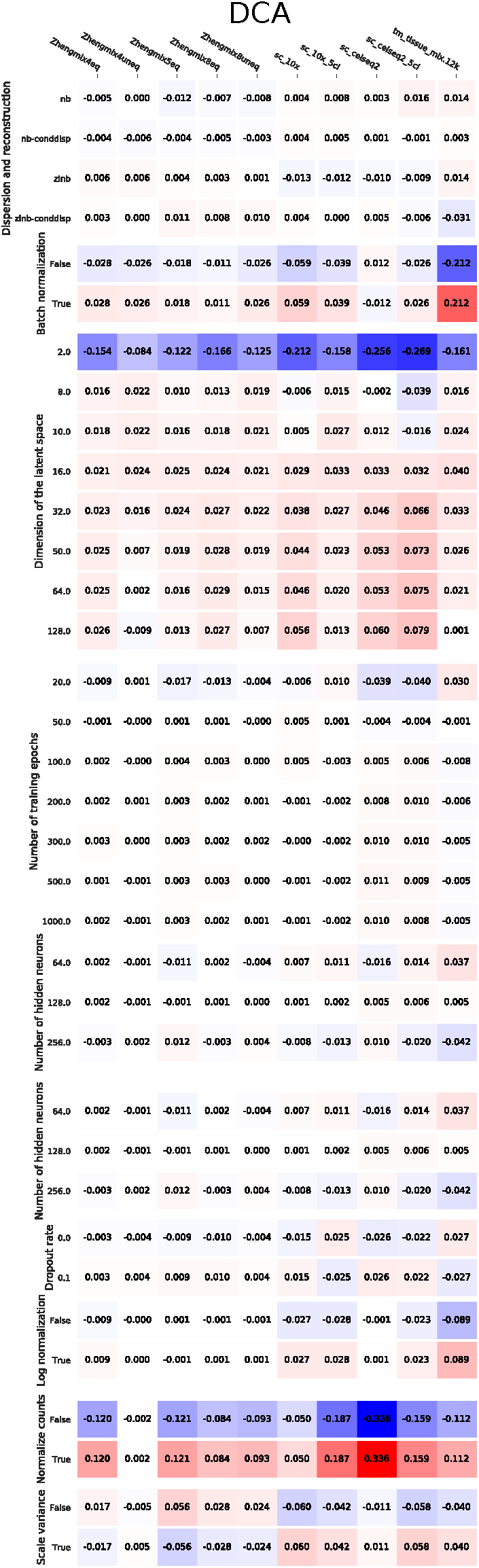
Heatmap visualization of the mean effect of each parameter value of DCA on its AMI. Each columns corresponds to a dataset. The rows are split by parameter and their values, the numbers show the average effect of that parameter value on the AMI compared to the mean AMI for DCA on that dataset. These effects come from a factorial ANOVA.

**Figure S13:**
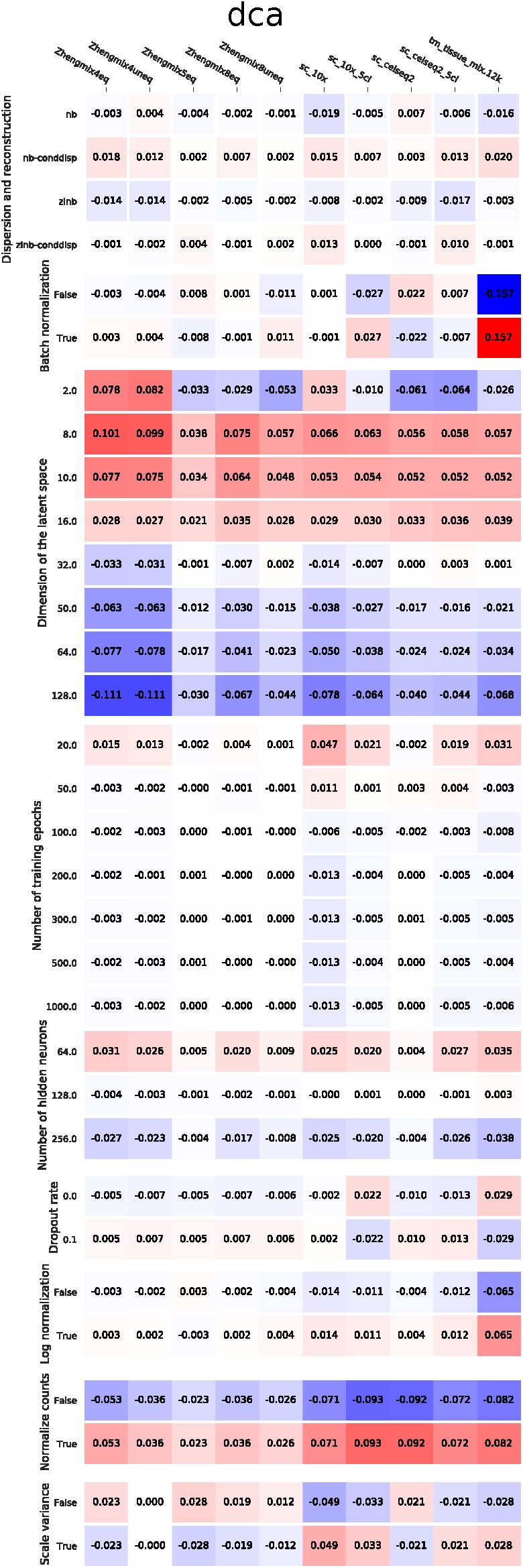
Heatmap visualization of the mean effect of each parameter value of DCA on its silhouette. Each columns corresponds to a dataset. The rows are split by parameter and their values, the numbers show the average effect of that parameter value on the silhouette compared to the mean silhouette for DCA on that dataset. These effects come from a factorial ANOVA.

**Figure S14:**
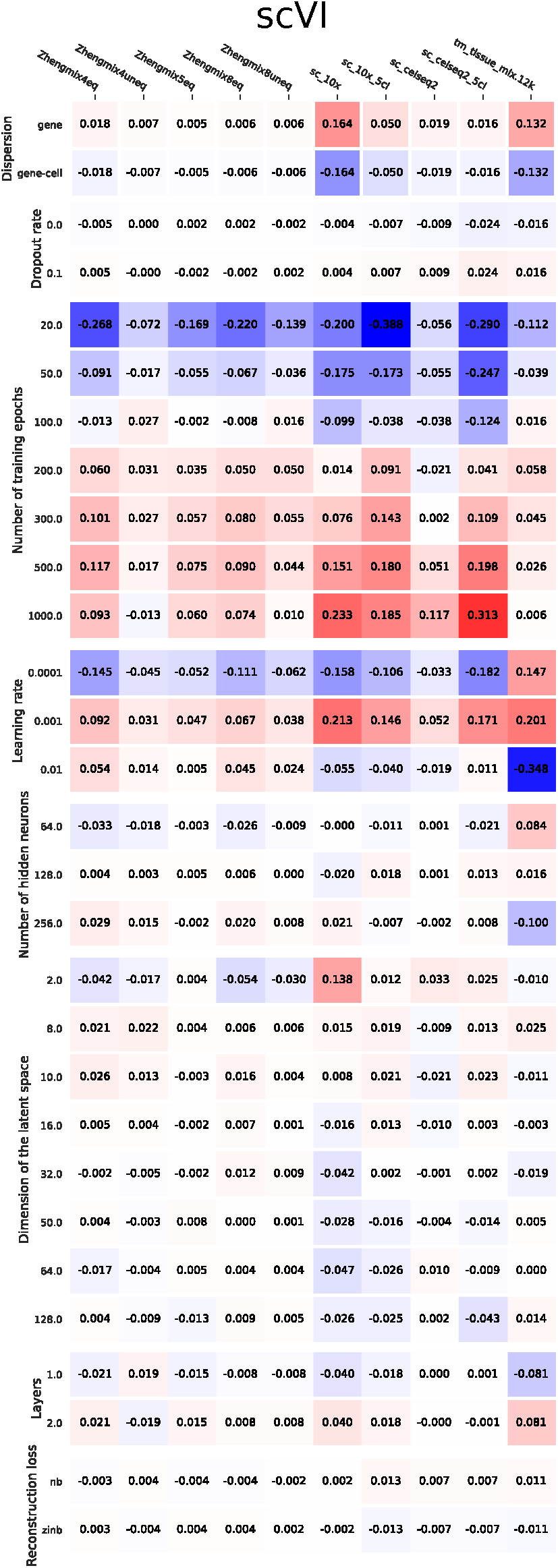
Heatmap visualization of the mean effect of each parameter value of scVI on its AMi. Each columns corresponds to a dataset. The rows are split by parameter and their values, the numbers show the average effect of that parameter value on the AMI compared to the mean AMI for scVI on that dataset. These effects come from a factorial ANOVA.

**Figure S15:**
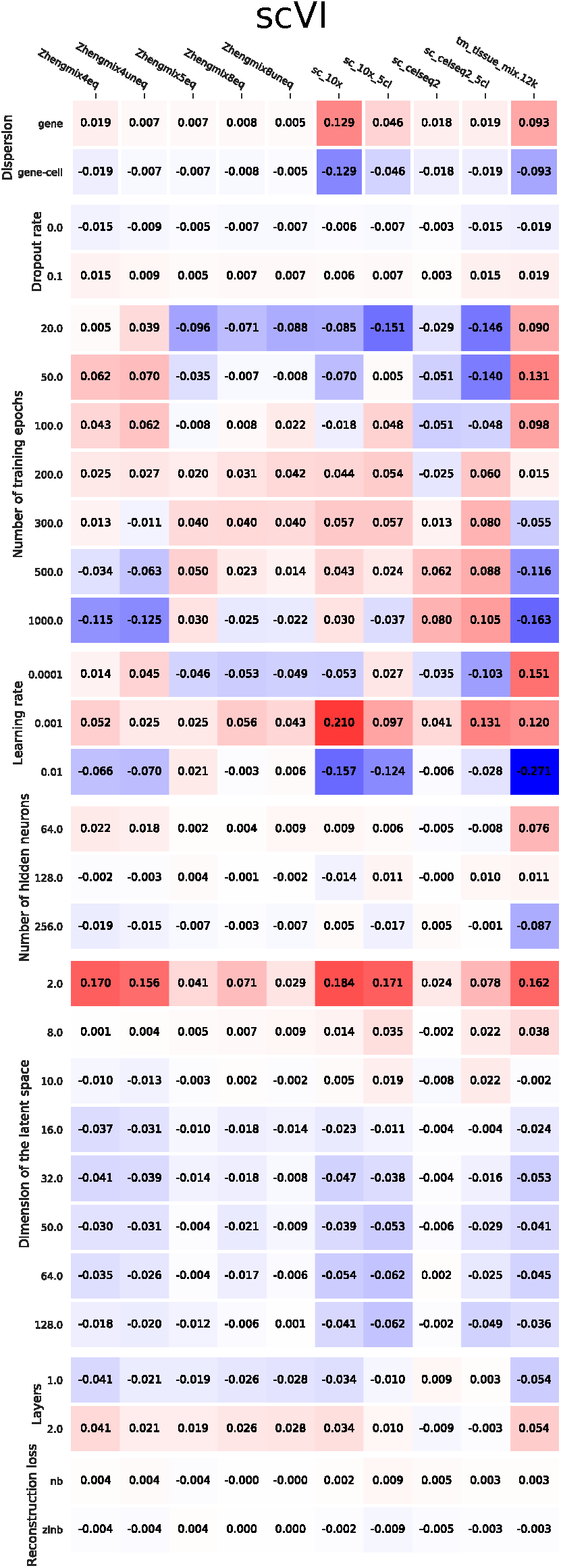
Heatmap visualization of the mean effect of each parameter value of scVI on its silhouette. Each columns corresponds to a dataset. The rows are split by parameter and their values, the numbers show the average effect of that parameter value on the silhouette compared to the mean silhouette for scVI on that dataset. These effects come from a factorial ANOVA.

**Figure S16:**
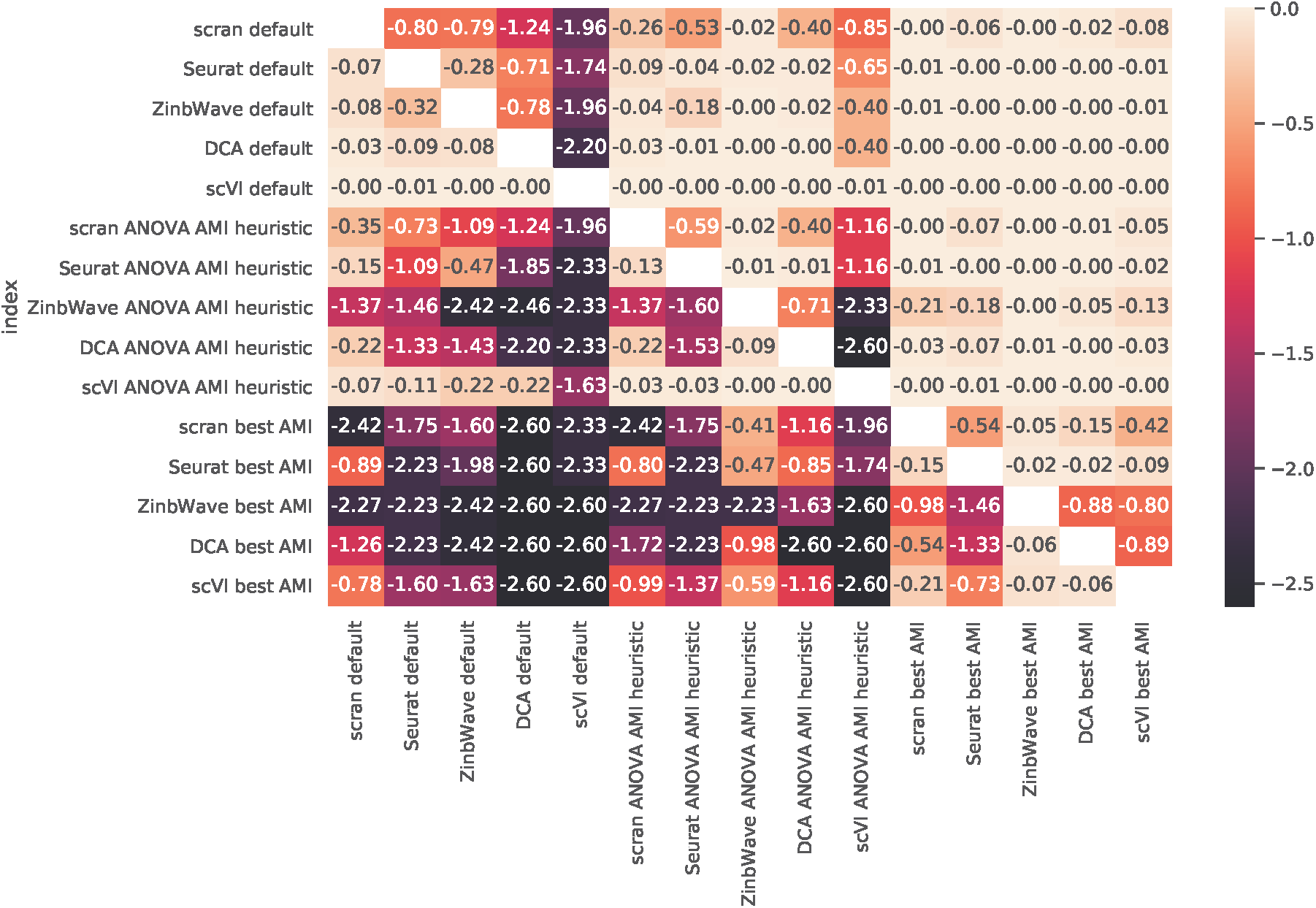
log base 10 p-values for the wilcoxon one-way test between the various methods and parameter configurations in AMI. A p-value of 0.05 coresponds to -1.3 in log base 10. The test is used to see if the method and parameters in the row achieve a higher AMI than the one in the column.

**Figure S17:**
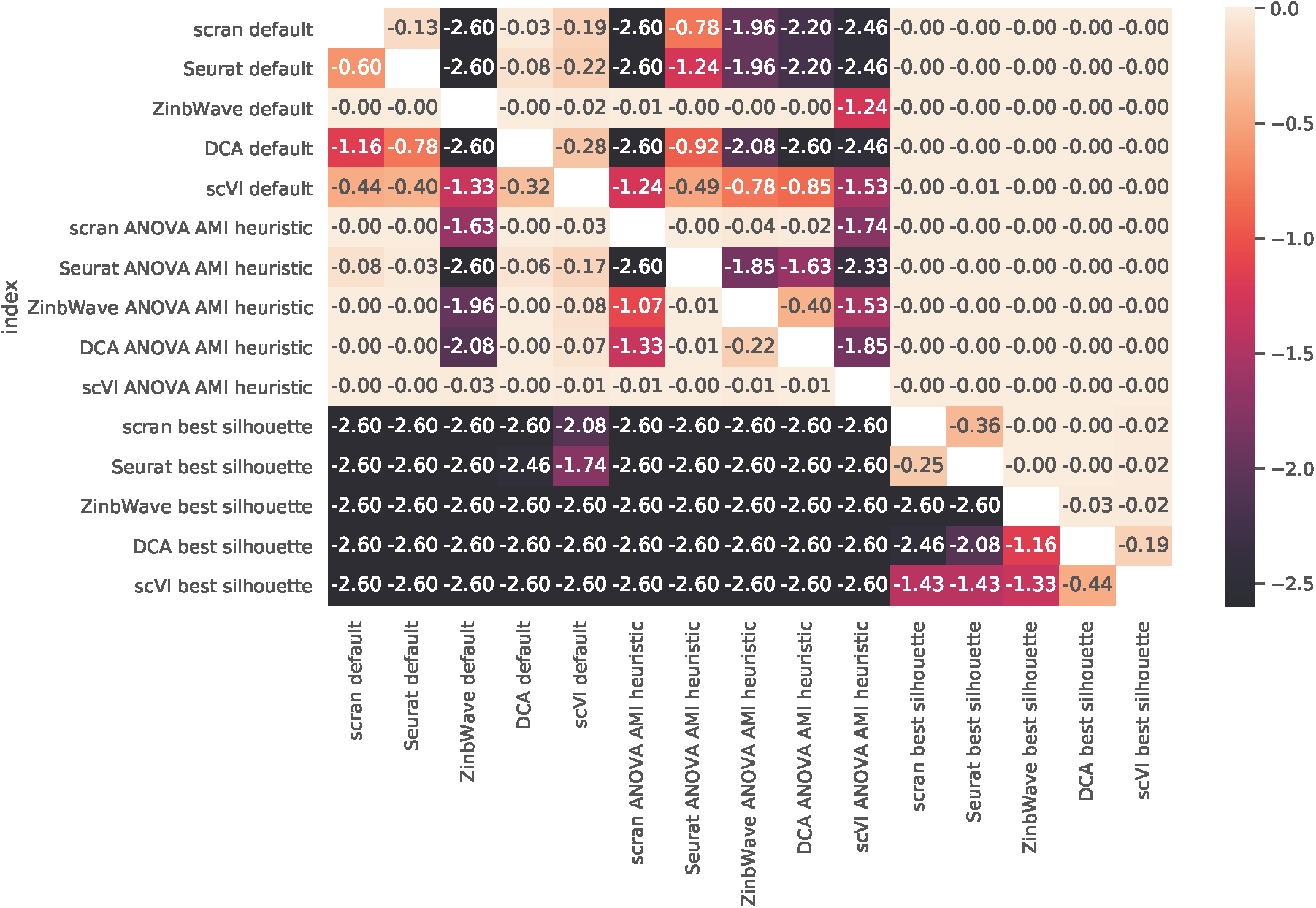
log base 10 p-values for the wilcoxon one-way test between the various methods and parameter configurations in silhouette. A p-value of 0.05 coresponds to -1.3 in log base 10. The test is used to see if the method and parameters in the row achieve a higher silhouette than the one in the column.

## Notes

https://doi.org/10.5281/zenodo.3966952

https://doi.org/10.5281/zenodo.3966234

### Competing Interest Statement

The authors have declared no competing interest.

### Summary of Updates

Revised version

